# Tree demographic strategies largely overlap across succession in Neotropical wet and dry forest communities

**DOI:** 10.1101/2023.06.14.544754

**Authors:** Markus E. Schorn, Stephan Kambach, Robin L. Chazdon, Dylan Craven, Caroline E. Farrior, Jorge A. Meave, Rodrigo Muñoz, Michiel van Breugel, Lucy Amissah, Frans Bongers, Bruno Hérault, Catarina C. Jakovac, Natalia Norden, Lourens Poorter, Masha T. van der Sande, Christian Wirth, Diego Delgado, Daisy H. Dent, Saara J. DeWalt, Juan M. Dupuy, Bryan Finegan, Jefferson S. Hall, José L. Hernández-Stefanoni, Omar R. Lopez, Nadja Rüger

## Abstract

Secondary tropical forests play an increasingly important role for carbon budgets and biodiversity conservation. Understanding successional trajectories is therefore imperative for guiding forest restoration and climate change mitigation efforts. Forest succession is driven by the demographic strategies – combinations of growth, mortality and recruitment rates – of the tree species in the community. However, our understanding of demographic diversity in tropical tree species stems almost exclusively from old-growth forests. Here, we assembled demographic information from repeated forest inventories along chronosequences in two wet (Costa Rica, Panama) and two dry (Mexico) Neotropical forests to assess whether the range of demographic strategies present in a community shifts across succession. We calculated demographic rates for >500 tree species while controlling for canopy status to compare demographic diversity in early successional (0-30 years), late successional (30-120 years) and old-growth forests. We quantified demographic diversity using two-dimensional hypervolumes of pairs of demographic rates and assessed whether shifts in demographic strategies were caused by intra-specific changes in demographic rates across succession or by species turnover. We expected that demographic strategies would shift from faster life-histories (fast growth, high mortality, high recruitment) in early successional forests to slower life histories (slow growth, low mortality, low recruitment) in old-growth forests and that shifts would be stronger in wet than in dry forests due to more pronounced differences in environmental conditions between early successional and old-growth forests. We also expected that demographic diversity would increase with succession. We found that demographic strategies largely overlapped across successional stages and that early successional stages already covered the full spectrum of demographic strategies found in old-growth forests. An exception was a group of species characterized by exceptionally high mortality rates that was confined to early successional stages in the two wet forests. Demographic diversity did not increase with succession. Our results suggest that current understanding of demographic strategies of tropical tree species, which has been generated mostly from long-term forest monitoring plots in old-growth forests, is largely representative of demographic diversity in general, and that demographic diversity recovers quickly during succession.

## INTRODUCTION

Tropical forests store almost half of global forest carbon and harbor a large proportion of the world’s biodiversity (Pan *et al*. 2011, Slik *et al*. 2015, FAO 2020). With only one third of tropical forests being undisturbed primary forests and rates of deforestation remaining high (Pan *et al*. 2011, FAO 2020), secondary tropical forests regrowing after land abandonment are of increasing importance for carbon storage and sequestration as well as biodiversity conservation (Chazdon *et al*. 2016, Arroyo-Rodríguez *et al*. 2017, Lewis *et al*. 2019, Rozendaal *et al*. 2019). Understanding successional trajectories is therefore imperative for guiding efforts of forest management and global change mitigation. Amongst a variety of factors, successional dynamics in a community are driven by the demographic strategies (or life-history strategies) of the component tree species (*sensu* Finegan 1996). However, empirical knowledge of how community-wide variation in demographic strategies changes along successional gradients remains limited.

Demographic strategies emerge from trade-offs that all organisms are faced with when allocating limited resources between fast growth, high survival or reproductive success (Stearns 1992, Metcalf & Pavard 2007) and that constrain the range of viable combinations of these demographic rates (Salguero-Gómez *et al*. 2016, Rüger *et al*. 2018). Recently, comparative analyses of life-history variation have improved our understanding of the consistency of demographic trade-offs structuring tropical forest communities (Kambach *et al*. 2022, Russo *et al*. 2021). However, most of our knowledge on demographic strategies stems from old-growth forests and it remains unknown how demographic diversity (i.e., community-wide variation in demographic strategies) in secondary forests compares to old-growth forests. Specifically, it is unclear whether certain demographic strategies are confined to certain successional stages.

As an example, Finegan (1996) describes Neotropical forest succession as the consecutive replacement of species with different life-history strategies, where early successional pioneer species will dominate the very first decades of succession but disappear later in succession. While we know that some early successional species can also occur in treefall gaps in old-growth forests (Schnitzer & Carson 2001), it is unclear whether there are species with unique demographic strategies that rely on large-scale disturbances for regeneration and therefore occur uniquely in early successional forests. If so, they might remain undetected when focussing tree demographic research exclusively on old-growth forests and important ecological processes might be overlooked.

In tropical wet forests, early successional environments are characterized by high resource levels (especially in terms of light availability; Montgomery & Chazdon 2001). Species that occur abundantly in these environments are, thus, commonly thought to trade off high growth and recruitment for high mortality rates (Tilman 1988, Finegan 1996, Chazdon 2014). These fast demographic rates are related to acquisitive values of functional traits such as low wood density or high specific leaf area and leaf nutrient content (Wright *et al*. 2010, Rüger *et al*. 2012, Rüger *et al*. 2018). On the other hand, species that are more abundant in late successional environments are expected to grow slowly but invest more resources in the ability to live for a long time, represented by trait values indicative of resource conservation such as high wood density or low specific leaf area (Tilman 1988, Finegan 1996, Chazdon 2014).

Indeed, using functional traits as proxies, several studies have confirmed that, in tropical wet forests, tree species with acquisitive trait values tend to dominate in early successional stages, while species with more conservative trait values gain in dominance as succession proceeds (e.g. Poorter *et al*. 2004, Dent *et al*. 2013, Lohbeck *et al*. 2013, Becknell & Powers 2014, Boukili & Chazdon 2017). Under such a scenario, we expect that the ranges of viable demographic strategies gradually shift from acquisitive strategies (fast growth, high mortality, high recruitment) towards more conservative strategies (slow growth, low mortality, low recruitment) during succession in tropical wet forests ( **Figure 1A**).

**Figure 1:**
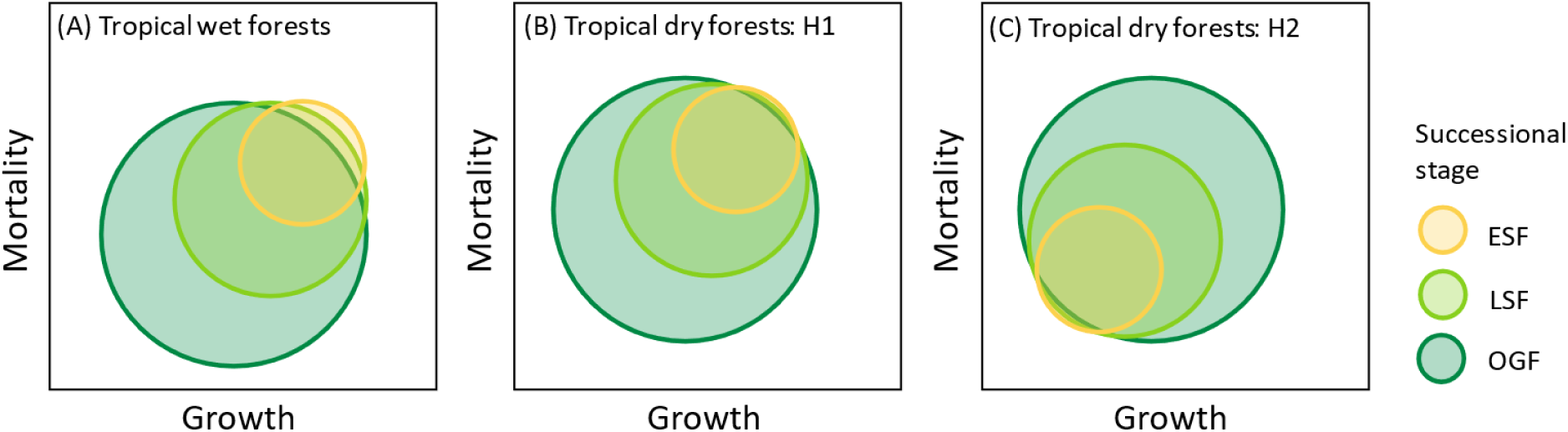
Hypotheses for potential shifts of the ranges of demographic strategies from early successional forests (ESF) to late successional forests (LSF) and old-growth forests (OGF) for (A) tropical wet forests and (B, C) tropical dry forests. H1 (B) and H2 (C) represent opposing hypotheses for tropical dry forests. It is not clear whether shifts from acquisitive to conservative demographic strategies or vice versa are to be expected since wood density and leaf dry matter content have been found to shift from conservative to acquisitive values, whereas specific leaf area has been found to shift from acquisitive to conservative values. Axis labels can be any combination of growth, mortality or recruitment since e.g. high values for all three demographic rates are expected in wet ESF.

In tropical dry forests, where water is considered a more important resource in shaping forest communities than light availability, early successional stages are characterized by dry and hot conditions changing towards moister and cooler environments as succession proceeds (Lebrija-Trejos *et al*. 2011, Pineda-García *et al*. 2013). Therefore, trait values have been found to reflect a shift from strategies associated with greater resource conservation and drought tolerance early in succession to more acquisitive strategies (e.g. lower wood density, leaf dry matter content) later in succession (Lebrija-Trejos *et al*. 2010, Lohbeck *et al*. 2013, Buzzard *et al*. 2016, Derroire *et al*. 2018, Poorter *et al*. 2019). However, acquisitive leaf trait values related to light capture efficiency (e.g. high specific leaf area) have also been found to decrease during succession (Lohbeck *et al*. 2013, Derroire *et al*. 2018), making unclear whether successional shifts towards more conservative (H1) or more acquisitive (H2) demographic strategies should be expected in tropical dry forests ( **Figure 1B & C**).

Environmental differences between early and late successional tropical dry forests are thought to be less pronounced compared to wet forests mainly due to greater canopy openness (Ewel 1977, Lebrija-Trejos *et al*. 2011). Indeed, Letcher *et al*. (2015) have observed a trend towards less successional habitat specialization among tree species in certain tropical dry forests. Based on this, we hypothesize that potential shifts in demographic strategies may be less pronounced in dry than in wet forests. However, as the horizontal and vertical heterogeneity of light availability increases across succession in both forest types, we hypothesize that the range of viable demographic strategies extends as succession progresses ( **Figure 1**).

Here, we assemble a unique chronosequence dataset of repeated forest inventories from four Neotropical forests varying in rainfall. We calculate demographic rates for >500 tree species and use hypervolumes to quantify demographic diversity in three successional stages. We address the following questions: (a) Do ranges of demographic strategies shift across succession in wet and dry tropical forests? (b) If so, are these shifts due to intra-specific changes in demographic rates across succession or due to species turnover? (c) Does demographic diversity increase with succession? Answers to these questions will reveal to what degree our understanding of demographic diversity gained from old-growth forests can be extended to secondary forests, for which much less information on demographic rates and strategies is available. This information will enhance our understanding of underlying mechanisms and improve our ability to predict successional dynamics in tropical forests with the help of demographic forest models.

## MATERIALS AND METHODS

### Study sites and forest inventory data

We used inventory data from nine long-term forest monitoring projects along chronosequences from four Neotropical lowland forest sites located in Costa Rica, Panama and Mexico. The sites differ in rainfall with mean annual precipitation ranging from 3,900 mm without any dry season to 900 mm with 90 % of annual rainfall occurring within 5.5 months of the year (**Table 1**). The forest in Costa Rica is a tropical wet evergreen broadleaved forest with a high proportion of palms (Clark & Clark 2000, Chazdon *et al*. 2007, Letcher & Chazdon 2009). The forest in Panama is a semideciduous tropical moist forest with a 3-month dry season (Denslow & Guzman 2000, van Breugel *et al*. 2013, Condit *et al*. 2019). The predominant natural disturbance regime in both wet sites are occasional windthrows and lightning strikes. The two sites in Mexico are both deciduous tropical dry forest differing from the wet sites in shorter stature, higher canopy openness and lower species richness (**Table 1**; Letcher *et al*. 2015). Forests in the Yucatán peninsula have undergone anthropogenic influences since ancient Mayan times and experience regular strong wind storms (Rico-Gray & García-Franco 1991, Hernández-Stefanoni *et al*. 2014, Saenz-Pedroza *et al*. 2020). The forest in Oaxaca is shorter in height, has arborescent cacti and has been only mildly affected by human disturbances (Lebrija-Trejos *et al*. 2008, Pérez-García *et al*. 2010, Gallardo-Cruz *et al*. 2010).

**Table 1:** Location, mean annual temperature (MAT), mean annual precipitation (MAP), dry season length (<100mm precipitation per month), number of species, number of species with 10 or more individuals, average old-growth forest (OGF) canopy height and length of the chronosequences used in this study. Note that the sampling area differs strongly between sites and the number of species included in the analyses may not be indicative of total species richness.

|  | Location | MAT (°C) | MAP (mm) | Length of dry season (months) | Number of species (N ≥ 10) | OGF canopy height* (m) | Chronosequence length |
| --- | --- | --- | --- | --- | --- | --- | --- |
| **Costa Rica** | 10°26' N, 84°00' W | 26 | 3.900 | - | 485 (355) | 20-35 | 1-57 years + OGF |
| **Panama** | 9°90' N, 79°51' W | 27 | 2.600 | 3 | 470 (391) | 15-28 | 0-120 years + OGF |
| **Yucatán, Mexico** | 20°05' N, 89°29' W | 26 | 1.100 | 6 | 154 (106) | 8-13 | 3-85 years |
| **Oaxaca, Mexico** | 16°39' N, 95°00' W | 28 | 900 | 7 | 125 (90) | 7-8 | 4-70 years + OGF |

\*Canopy heights are from Clark *et al.* (2021), Mascaro *et al.* (2011), Dupuy *et al.* (2012) and Lebrija-Trejos *et al.* (2008).

The four chronosequence sites comprised a total of 252 secondary forest plots and 23 old-growth forest plots ranging in size from 0.04 ha to 50 ha (Table S1, Figure S1). Plots are located in complex landscapes mainly consisting of fragments of old-growth and second-growth forest, plantations, agricultural land and pastures. Most secondary forest plots were established on abandoned agricultural land used primarily for low-intensity crop farming or cattle ranching (Denslow & Guzman 2000, Chazdon *et al*. 2007, Letcher & Chazdon 2009, Lebrija-Trejos *et al*. 2011, van Breugel *et al*. 2013). Some plots were only clear cut but not farmed (Chazdon *et al*. 2007, Letcher & Chazdon 2009). In general, previous forest vegetation was completely removed, yet in a few cases some remnant trees remained, which we excluded from the analyses. The age of the youngest plots ranged from 0 to 4 years across sites, whereas the oldest secondary forest plots ranged from 57 to 120 years after agricultural abandonment (Figure S2). All free-standing woody individuals above the plot-specific size threshold (generally 1 or 5 cm diameter at breast height (dbh); range: 1-10 cm dbh, Table S1) were measured, marked and remeasured 1 to 10 years later. We selected census intervals of 5 years, if possible (range: 4-10 years, Table S1). In the wet sites (Costa Rica, Panama), only the largest stem of an individual was measured in some plots. In the dry sites (Yucatán, Oaxaca), where resprouting is an important mode of regeneration and, thus, multi-stemmed individuals are abundant (Vieira & Scariot 2006), all stems of an individual were measured, but not individually marked.

We assigned all census intervals to one of three successional stages (Figure S2). Census intervals from secondary forest plots ending less than 30 years after abandonment were classified as early successional forests (ESF) and intervals ending less than 120 years after abandonment were classified as late successional forests (LSF), although most plots were not older than 90 years. Census intervals from old-growth forest plots were classified as old-growth forests (OGF). Data from old-growth forests in Yucatán was not available to sufficient extent.

### Canopy layer assignment

Growth and mortality rates of individual trees depend on their size and light availability. To account for these differences, we assigned trees to discrete canopy layers based on their size and the size of their neighbors following the Perfect Plasticity Approximation approach of Purves *et al*. (2008) and Bohlman & Pacala (2012). To do this, we first divided all plots into subplots that were either predefined by the sampling design, or trees were assigned to subplots based on their spatial coordinates. The size of these subplots ranged from 625-1000 m² in wet sites (except for a 100 m² plot in a <20 year old forest) depending on the sampling design and plot sizes (Table S1). In the dry sites, where trees are generally smaller than in wet forests, subplot sizes ranged from 100-125 m² except for some 400 m² plots in Yucatán (Table S1). Next, we sorted trees by dbh of the largest living stem within subplots. We then estimated the crown area for all trees using allometric equations (see SI Methods). Starting from the largest, we assigned trees to the top canopy layer (layer 1) as long as the cumulative estimated area of their crowns did not exceed the subplot area. Smaller trees were successively assigned to lower canopy layers in the same way. Calculating demographic rates in discrete canopy layers has proven useful in capturing variance in demographic strategies between co-occurring species (Bohlman & Pacala 2012, Rüger *et al*. 2018) and in predicting forest dynamics (Rüger *et al*. 2020).

### Demographic rates

We calculated demographic rates for species with interpretable stem growth, i.e. excluding palms and hemi-epiphytes. We determined dbh increment and mortality for all observations and subsequently calculated species-level annual growth and mortality rates for each canopy layer and successional stage. Individual annual tree growth *g^i^* was calculated as

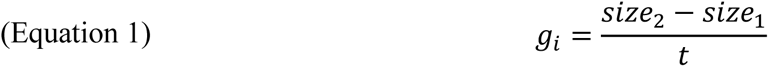

with *size* being the dbh of the largest living stem of an individual in wet sites and dbh equivalent of the total basal area (ba) of all living stems of an individual in dry sites in the first and second census, respectively, and *t* being the time elapsed between the two size measurements in years. We used dbh equivalent of the total basal area as the measure of size because in the dry sites, stems were not individually marked and, thus, stem-level dbh growth could not be calculated. Species-level growth rates per canopy layer (*g_j,l_*) were calculated as the median growth of all individuals *i* of species *j* in layer *l*:

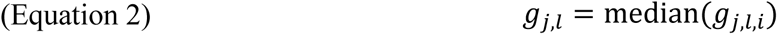

Species-level annual mortality rates per canopy layer (*m_j,l_*) were determined as

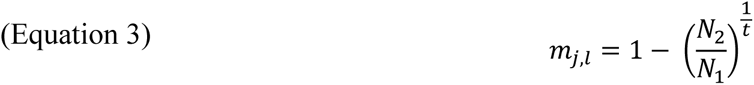

with *N_1_* being the number of living individuals in the first census, *N_2_* being the number of individuals remaining alive in the second census and *t* being the mean census interval length in years (measured to the nearest day). Multi-stemmed individuals were deemed alive if at least one stem was alive and dead if all stems were dead.

Species-level per-capita recruitment rates for each successional stage were determined as

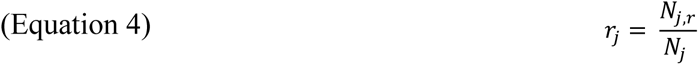

with *N_j,r_* being the annual number of recruits per hectare that surpassed the 1 cm dbh threshold between two consecutive censuses divided by the mean census interval length in years and *N_j_* being the average number of individuals per hectare of the respective species in the respective successional stage across plots. Only plots or subplots with a minimum dbh threshold of 1 cm were used to determine *N_j,r_*, whereas all plots of the respective successional stage were used to determine *N_j_*. In some successional stages, only a few small plots had a minimum dbh threshold of 1 cm, which limited our ability to assess recruitment rates. However, recruitment over the 1 cm dbh threshold is a more frequent event than recruitment over the 5 cm dbh threshold. Thus, recruitment rates could be quantified for more species using the lower threshold.

### Quantification of demographic diversity

We used two-dimensional hypervolumes based on gaussian kernel density estimations (Blonder *et al*. 2014) to represent and quantify diversity of demographic strategies spanning pairs of demographic rates: growth vs mortality, growth vs recruitment and mortality vs recruitment. Because many species did not occur in all canopy layers and species with non-observed values cannot be included in hypervolume analyses, we calculated hypervolumes using only growth and mortality rates from a single canopy layer. Only species with at least 10 or 5 observations for mortality per canopy layer were included in the analyses for wet and dry sites, respectively. Species with mortality rates of 0 or 1 or with recruitment rates of 0 were excluded from the respective hypervolumes. A total of 353, 503 and 463 species met the criteria for inclusion in the analyses in canopy layers 1, 2 and 3, respectively. We used growth and mortality rates from canopy layer 2 to maximize the number of species included in the analyses (Tables S3-S5). To ensure representability, we also examined hypervolumes using growth and mortality rates from canopy layers 1 and 3.

We natural log transformed all demographic rates to ensure approximate gaussian multivariate distributions. We estimated kernel bandwidths (the parameter defining the smoothness of the probability densities) for each hypervolume per site using the cross-validation method (Duong 2007, Blonder *et al*. 2014). We used the same bandwidths for all successional stages per site to ensure comparability among sites (Blonder 2018). Hypervolume boundaries represent the smallest volume that captures 80% of the total probability densities. All species contributed equally to the hypervolume calculations. We used the *hypervolume* R package (version 3.0.4) for all analyses (Blonder *et al*. 2014). We also report demographic spaces with underlying abundance heatmaps to indicate observed shifts in the dominance of certain demographic strategies.

To quantify overlap of demographic strategies in different successional stages, we calculated overlap statistics for all hypervolumes using the *hypervolume_overlap_statistics* function. We used bootstrapping and rarefaction techniques (r = 100 replicates, n = 10 species per replicate) to account for differences in the number of species included and to obtain 95% confidence intervals. Likewise, we derived volumes of all two-dimensional hypervolumes (i.e. areas) to quantify the amount of demographic diversity using the *hypervolume_gaussian* function. We assumed statistically significant differences if the confidence intervals did not overlap.

To evaluate whether shifts in the range of demographic strategies were due to intra-specific variation in demographic rates across successional stages, we performed major axis regressions between species’ demographic rates in different successional stages. To evaluate whether shifts in the range of demographic strategies were due to species turnover, we assessed successional trends in the abundance of species that exhibited a demographic strategy that was exclusive to one particular successional stage. As we only found unique demographic strategies in the early successional stage, we modeled abundances per ha of the species with this unique strategy as a function of stand age with the form: Ln(abundance) = a * stand age^b^. Parameters a and b were estimated using the *nls* R function. For parameter estimation, we only used data points from the point in time when species reached their highest abundance onwards.

All analyses were carried out in R version 4.2.2 (R Core Team 2022). Taxonomy was standardized according to The Plant List version 1.1 (http://www.theplantlist.org) using the *Taxonstand* package (Cayuela *et al*. 2012).

## RESULTS

We used 1,385,018 observations from 352,243 individual trees to calculate demographic rates for a total number of 503 species from 77 families. The number of individual trees ranged from 3,302 in Oaxaca to 312,328 in Panama (Table S2).

### Overlap in demographic strategies across succession

In all forest sites, the ranges of demographic strategies present in the three successional stages largely overlapped (**Figure 2**), except for recruitment rates that shifted slightly towards fewer recruits during succession in all sites except Yucatán. In the wet sites, we found a demographic strategy exclusive to early successional forests, which was primarily associated with exceptionally high mortality rates of 10% or more (**Figure 2A & B**). This group of high mortality species consisted of 17 and 28 species in Costa Rica and Panama, respectively. Many of these species are typically considered pioneer species (e.g. *Cecropia insignis, Ochroma pyramidale, Trema integerrima, Byrsonima crassifolia, Conostegia xalapensis, Vernonanthura patens, Vismia baccifera, Vismia macrophylla*; see Figures S15 & S16). Many species within this group did not grow particularly fast, especially in Panama. In Costa Rica, no recruits were recorded for most of the species in this group (**Figure 2I**), whereas in Panama, most of these species had fairly high recruitment rates (**Figure 2J**).

**Figure 2:**
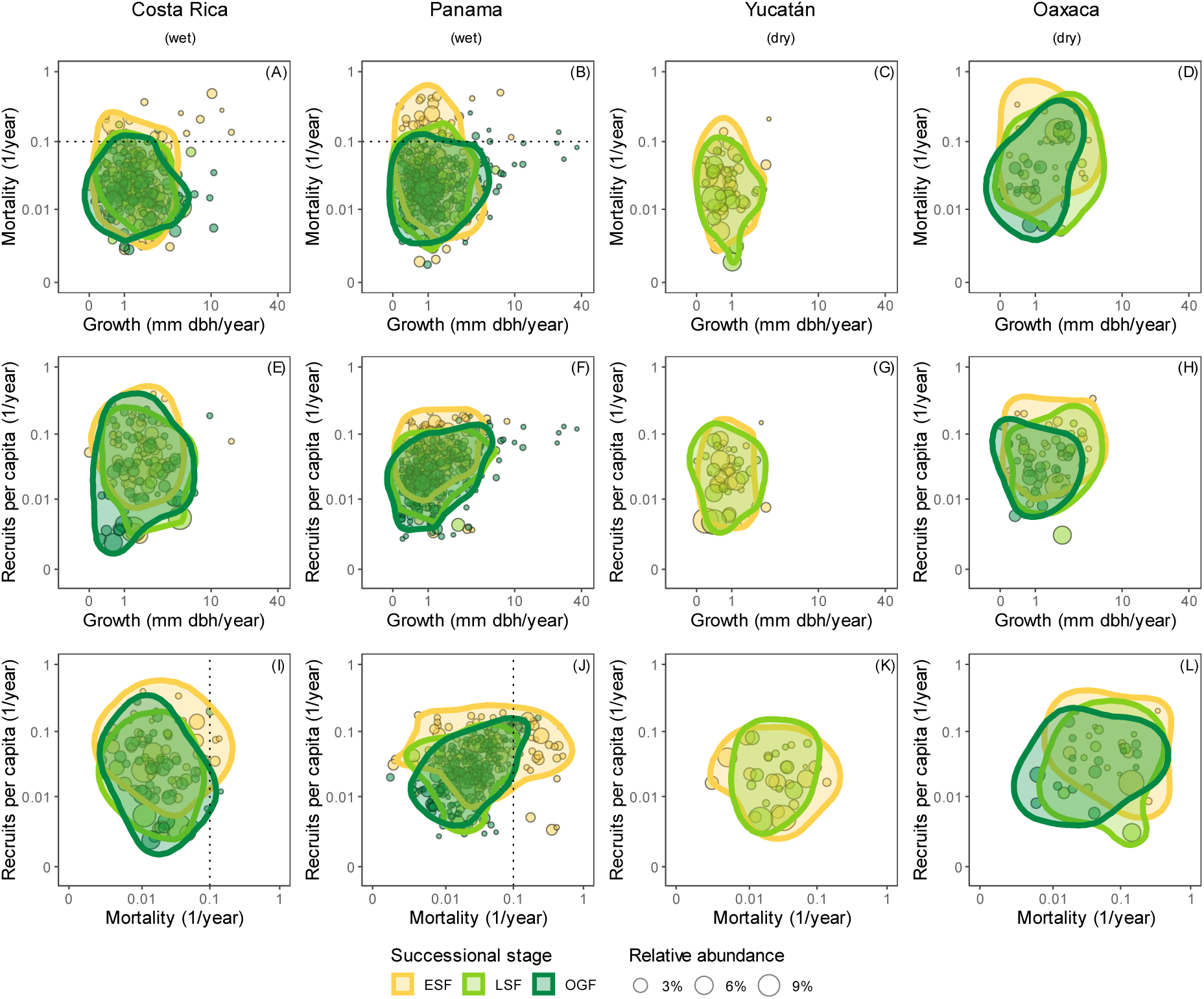
Two-dimensional hypervolumes representing ranges of demographic strategies for different pairs of demographic rates (A-D: growth-mortality, E-H: growth-recruitment, I-L: mortality-recruitment) for all sites and successional stages (ESF = early successional forest, LSF = late successional forest, OGF = old-growth forest). All axes are log-transformed (Ln). Growth and mortality rates are from individuals assigned to canopy layer 2 because this layer contains the most individuals and species. Hypervolume boundaries represent the smallest volume that captures 80% of the total gaussian probability densities. All species contributed equally to the hypervolume calculation. Points represent species and point sizes indicate relative abundances within the successional stage.

In the dry sites, we did not identify generalizable shifts of demographic strategies across succession. In Oaxaca, but not in Yucatán, the range of growth rates shifted slightly towards slower growth during succession. In Oaxaca, but not in Yucatán, the range of mortality rates shifted slightly towards lower values in older forests. The range of growth and mortality rates was smaller in Yucatán compared to Oaxaca.

These results were robust to the choice of canopy layer (Figures S3-S5) and were not biased by the number of species included (Figure S6). The measured average hypervolume overlap across pairs of demographic rates between successional stages did not differ across sites and was also similar among all successional stages within sites (**Figure 3 Figure 6**, Figure S7).

**Figure 3:**
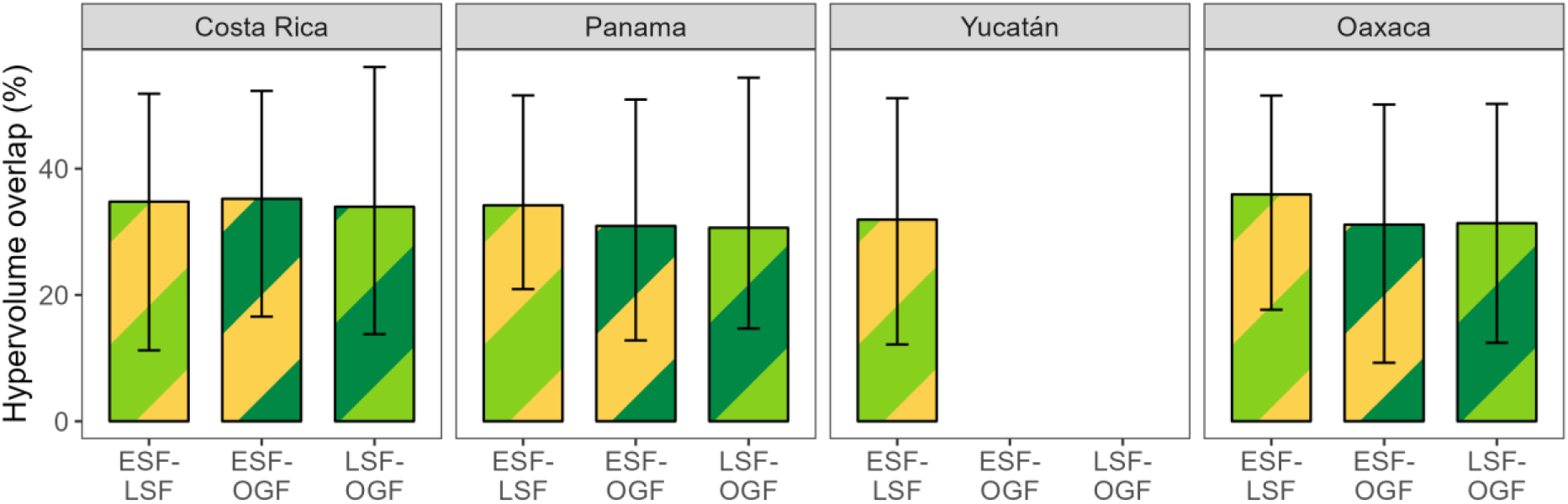
Mean overlap statistics of the two-dimensional hypervolumes representing the ranges of demographic strategies for all successional stages (ESF = early successional forest, LSF = late successional forest, OGF = old-growth forest). Colored bars represent the median rarefied and bootstrapped values, error bars represent 95% confidence intervals (r = 100 replicates, n = 10 species per successional stage). All values are means across pairs of demographic rates. Individual values per pair of demographic rates are given in Figure S7.

Except for Oaxaca, species abundance was more evenly distributed across the range of demographic strategies in early successional forests compared to later successional stages, where species with the highest abundances were more concentrated around conservative strategies (Figures S8-S11).

### Are shifts in demographic strategies due to intra-specific variation or due to species turnover?

Overall, intra-specific variation in demographic rates across succession was low in all sites (**Figure 4**, Figures S12-S14). Especially growth and mortality rates were generally consistent across successional stages. In Panama and Oaxaca, recruitment rates were higher in secondary forests than in old-growth forests (**Figure 4**, Figure S14), whereas in Costa Rica and Yucatán, results for the major axis regressions on recruitment rates were inconclusive (Figures S12 & S13).

**Figure 4:**
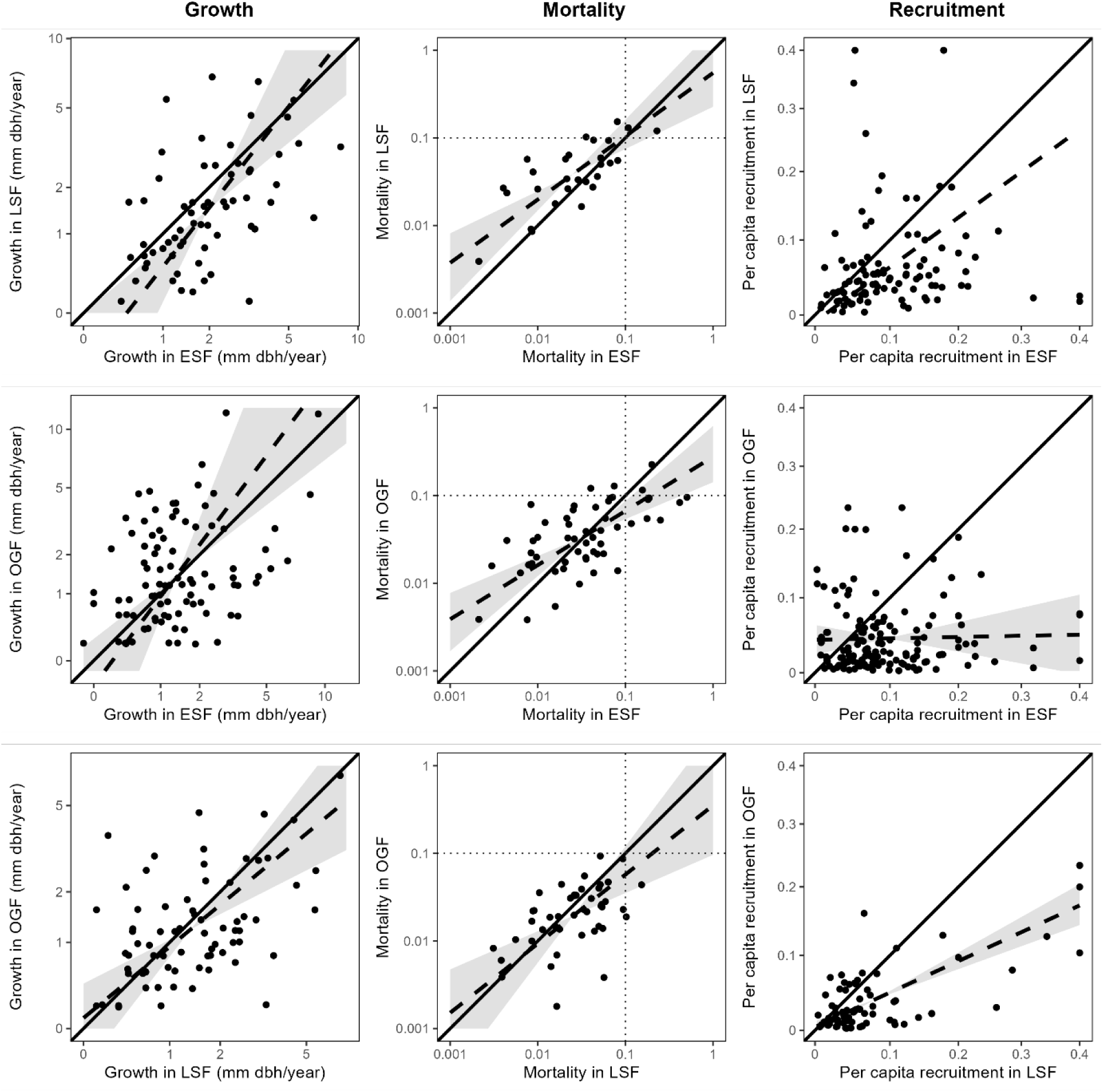
Major axis regressions for species’ demographic rates in early successional forests (ESF), late successional forests (LSF) and old-growth forests (OGF) in Panama (for other sites see Figures S12-S14). Each point represents a species and its demographic rates in the respective successional stage. Solid lines represent the 1:1-line, dashed lines represent the major axis regression lines and areas highlighted in grey represent the confidence intervals. Growth and mortality rates are from canopy layer 2. Only species with at least 5 observations for growth and survival in both successional stages were included, respectively. If no confidence intervals are given, the model was not statistically significant (i.e., the variables are unrelated, p ≥ 0.05).

Most of the species that exhibited an exclusive demographic strategy (i.e., species with mortality >10% in wet ESF) decreased substantially in abundance during the first 30 years of succession (**Figure 5**, Figures S15 & S16). In Costa Rica, only five out of the 17 species were also found in LSF (four species) or OGF (one species). In Panama, eleven out of the 28 species forming this group in ESF were also found in LSF (two species) or OGF (nine species), albeit at low abundances. Species from this group that were present in more than one successional stage generally had lower mortality rates in later successional stages (**Figure 4**, Figure S14).

**Figure 5:**
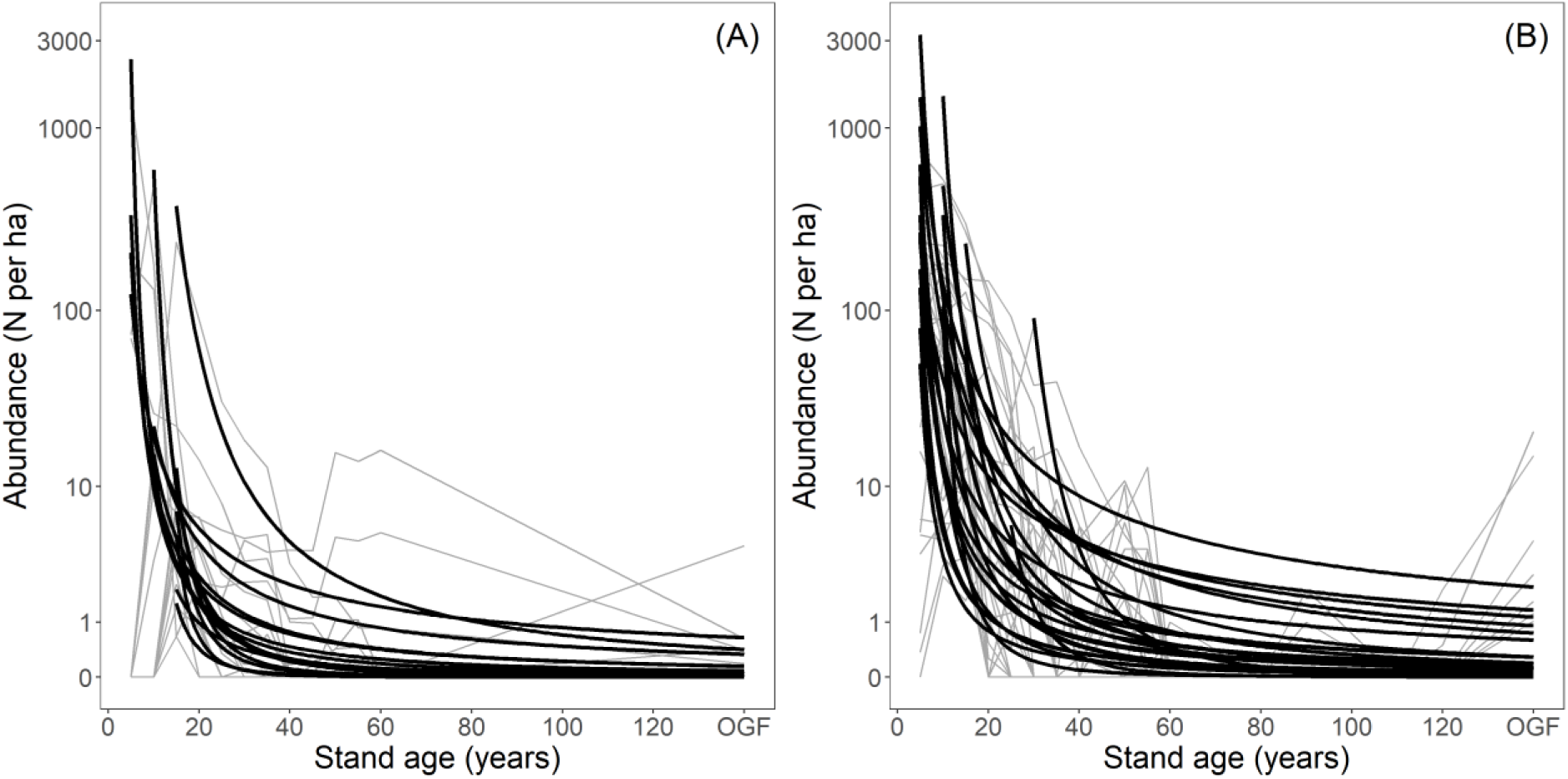
Observed (grey) and modelled (black) abundances > 1 cm dbh over time of species with annual mortality >10% exclusive to wet early successional forests in (A) Costa Rica and (B) Panama. Models are of the form Ln(abundance) = a*stand age^b. Parameters a and b were estimated using the *nls* R function. The model did not converge for two species in Costa Rica and three species in Panama due to irregular patterns in abundance over time. The model does not accurately capture that one species in Costa Rica and seven species in Panama had increased abundances in OGF. Individual models are shown in Figures S15 and S16.

### Demographic diversity does not increase with succession

We found that demographic diversity (quantified as the area of the 2-dimensional hypervolumes) did not increase with succession (**Figure 6**, Figure S17). In the wet sites, demographic diversity tended to decrease, but not significantly. In the dry sites, demographic diversity tended to increase, but not significantly.

**Figure 6:**
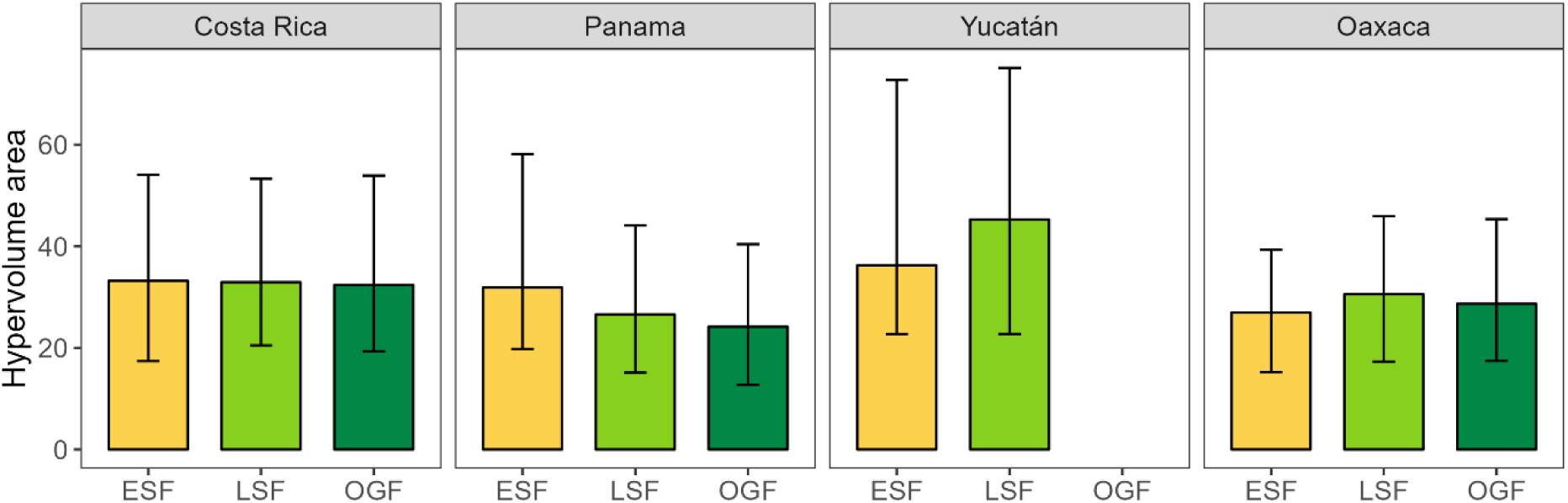
Mean areas of the two-dimensional hypervolumes representing the ranges of demographic strategies for all successional stages (ESF = early successional forest, LSF = late successional forest, OGF = old-growth forest). Colored bars represent the median rarefied and bootstrapped values, error bars represent 95% confidence intervals (r = 100 replicates, n = 10 species per successional stage). All values are means across pairs of demographic rates. Individual values per pair of demographic rates are given in Figure S17.

## DISCUSSION

We used demographic rates from 503 woody plant species to compare ranges of demographic strategies along successional gradients in four Neotropical forests. Contrary to our expectations, we found not only that demographic strategies largely overlapped across successional stages, but also that the amount of demographic diversity was similar along succession. Interestingly, we found a group of species with exceptionally high mortality rates that occurred exclusively in early successional forests in wet sites. Our results suggest that insights gained from analyses of demographic rates in old-growth forests are largely representative for forests of all successional stages, with the exception of a strategy associated with high mortality that, in wet forests, only occurs in early successional forests.

### Demographic strategies largely overlap across succession

In contrast to our expectations, we found large amounts of overlap of demographic strategies across successional stages in all four sites. Almost all demographic strategies that were present in old-growth forests were present in secondary forests after 30 years of succession in both wet and dry tropical forests. This could suggest that most species, regardless of their life-history strategies, can establish in early successional forests as long as their seeds reach the site, and highlight the importance of stochastic processes in general and of dispersal limitation in particular for successional trajectories (Chazdon 2008, Norden *et al*. 2015, Dent & Estrada-Villegas 2021). At the same time, tropical wet forests recover quickly and the range of microsites that can develop during the first 30 years of succession may accommodate the full range of demographic strategies exhibited by species that occur in older forests. This does not mean that abundances of species with different demographic strategies do not shift across succession (Rüger *et al*. 2022), but here we focus on the presence or absence of demographic strategies.

We found a shift towards lower recruitment rates during succession in all sites except Yucatán. Recruitment rates strongly depend on seedling performance and, hence, on resource (primarily light) availability at the forest floor (Montgomery & Chazdon 2001, Kitajima *et al*. 2013, Kupers *et al*. 2019). Because light availability at the forest floor decreases during succession (Denslow & Guzman 2000), recruitment rates are predicted to do the same.

In Oaxaca, the driest forest site, the range of demographic strategies shifted slightly towards lower growth, mortality and recruitment rates, i.e., towards more conservative life-history strategies during succession. In Yucatán, the second dry forest site, however, we detected a slight shift towards higher recruitment, hindering our ability to generalize more broadly from our results. The forest in Yucatán stands out in that it occurs in a landscape that has been shaped by human land use for many centuries. Thus, the pool of tree species might have been restricted over time to those species that are able to cope with frequent disturbance, including the ability to resprout (Rico-Gray & García-Franco 1991, Kammesheidt 1999, Sanaphre-Villanueva *et al*. 2017). This is also indicated by the smaller range of growth and mortality rates of the dry forest in Yucatán compared to that of Oaxaca.

### Species with high mortality rates are exclusive to early succession in wet forests

We expected to find the most acquisitive demographic strategies with highest growth, mortality and recruitment rates in early successional wet forests. Yet, the group of species exclusively observed in these forests was associated with high mortality and moderately high recruitment rates, but not particularly with fast growth. Potentially, higher growth rates might be masked because the entire lifecycle of these short-lived species is completed within the early successional stage (0-30 years since abandonment), including senescent stages when growth might decline.

Although some of these high-mortality species were highly abundant in early successional forests, no recruits were recorded for most of them in Costa Rica. Here, recruitment rates might be less informative than in other forests because only a few plots that were 12 years or older had information on trees ≥ 1 cm dbh and met our criterion for the calculation of recruitment rates (Table S1). Additionally, recruitment in the plots at the La Selva Biological Station (referred to in Table S1 as Sarapiquí) is known to be affected by collared peccaries (Kuprewicz 2013). Given their high abundance during the first ∼15 years of succession, many of the high-mortality species might actually have similarly high recruitment rates in early successional forests as many of the high-mortality species in Panama, where data availability was more consistent throughout the chronosequence. Hence, in contrast to common assumptions, early successional specialist demographic strategies might trade off high mortality for high recruitment rather than consistently high growth rates.

In tropical dry forests, species that are present in early successional forests can persist for a longer time and do not have a unique demographic strategy. Because of a lower and (seasonally) more open canopy, early and late successional environments are less contrasting in dry compared to wet forests (Lebrija-Trejos *et al*. 2011, Letcher *et al*. 2015). Moreover, many resprouting species in early successional forests might in fact be species that were abundant pre-disturbance and therefore follow demographic strategies associated with late successional environments (Boucher *et al*. 2001, Lebrija-Trejos *et al*. 2008). Additionally, resprouting trees in early successional forests likely rely on belowground carbohydrate reserves of the old root system and therefore might have similar demographic rates as in old-growth forests (Poorter *et al*. 2010).

### Shifts in demographic strategies are mainly due to species turnover

The shifts that we identified mainly relate to the loss of a group of species with exceptionally high mortality rates (>10%), that is present only in early successional tropical wet forests. The majority of species within this group did not persist in later successional stages, suggesting that their disappearance is primarily due to species turnover as projected by Finegan (1996). However, the few species from this group that are present in old-growth forests do exhibit lower mortality rates there, indicating that both species turnover as well as intra-specific variation contribute to this process.

### Demographic diversity does not increase with succession

Counter to our expectations, we did not find a general pattern of increasing diversity in demographic strategies during succession. Indeed, demographic diversity seems to recover to old-growth forest values within the first 30 years of succession. Similarly, Poorter *et al*. (2021) found that structural heterogeneity and species richness in secondary tropical forests recovered to 90% of old-growth forest values at around 30 years after abandonment, whereas species composition only recovered after more than a century (Poorter *et al*. 2021). This suggests that demographic diversity is more closely linked to species richness than to species composition, indicating that many different species exhibit similar demographic strategies and fill similar demographic niches.

### Limitations

When interpreting our results, it should be considered that we use a chronosequence approach that substitutes space for time and thus infers temporal trends from static data (Foster & Tilman 2000, Johnson & Miyanishi 2008, Walker *et al*. 2010). Moreover, data availability as well as data collection methodologies varied widely across sites (Figure S1), and plots within each chronosequence also varied in extent and minimum dbh threshold (Table S1). Nevertheless, our results are robust to this heterogeneity and independent from the number of species included (e.g. Figure S6).

## CONCLUSION

Overall, we find that secondary forests harbor similar levels of demographic diversity as old-growth forests, indicating that early successional stages already contain the full spectrum of life-history strategies found in old-growth forests, and that demographic data from old-growth forests is surprisingly informative for understanding the diversity of demographic strategies in tropical forests in general. Our results also suggest that the recovery of demographic diversity is more closely linked to species richness than to species composition. Lastly, our results indicate that, contrary to common assumptions, early successional specialists in tropical wet forests trade off high mortality for high recruitment rates rather than consistently fast growth, at least when integrating over 30 years of successional development.

Our results enrich the current understanding of tropical secondary succession by using a demographic perspective that evaluates mechanisms that underpin succession. As ranges of demographic rates are similar and species-specific demographic rates are largely consistent across succession, we argue that demographic information from old-growth forests can be used to predict successional changes in the dominance of different species or species groups and to estimate future tropical forest carbon stocks with the help of demographic forest models (Purves *et al*. 2008, Rüger *et al*. 2020), especially for tropical dry forests. Accurate predictions of early successional dynamics in tropical wet forests, however, likely rely on information about demographic strategies that occur uniquely during the first 30 years of succession.

## ACKNOWLEDGEMENTS

This paper is a product of the sDiv working group sUCCESS. We thank the owners of the forest sites for access to their forests, all the people who have established and measured the plots, the institutions and funding agencies that supported them (see below). **Funding:** This research was supported by the German Centre for Integrative Biodiversity Research (iDiv) Halle-Jena-Leipzig (sDiv W7.20 sUCCESS to LP, NR, and MvB, iDiv-Flexpool grants 34600967 and 34600970 to NR) funded by the Deutsche Forschungsgemeinschaft (DFG; FZT-118); Netherlands Organisation for Scientific Research - NWO (ALW.OP241 to LP, MTvdS, and CCJ; ALW.OP457 to FB, RM, LP and JAM; and Veni.192.027 to MTvdS); Fundação de Amparo á Pesquisa do Estado de São Paulo (17418 NEWFOR to FB); Agencia Nacional de Investigación y Desarrollo (FONDECYT Regular No. 1201347 to DC); Conselho Nacional de Desenvolvimento Científico e Tecnológico (SinBiose-REGENERA 442371/2019-5 to CCJ); Fondo Mixto CONACYT - Gobierno del estado de Yucatán (FOMIX YUC-2008-C06-108863 to JMD and JLHS, FOSEMARNAT 2004-C01-227, Reinforcing REDD+ and the South-South Cooperation Project, CONAFOR and USFS to JLHS); STRI, ForestGEO, Heising–Simons Foundation, HSBC Climate Partnership, Stanley Motta, SmallWorld Institute Fund, the Hoch family (to JSH and MvB); Universidad Nacional Autónoma de México, Programa de Apoyo a Proyectos de Investigación e Innovación Tecnológica (DPAGA–PAPIIT IN218416, DPAGA– PAPIIT IN217620 to JAM and RM); SENACYT Panama Grant (COL10-052 to DHD, SJD and ORL); US National Science Foundation (DEB-9208031 to DHD and SJD and EAR-1360391 to MvB); Yale-NUS College and MOE (through a startup grant and grant IG16-LR004 to MvB). The BCI forest dynamics research project was founded by S.P. Hubbell and R.B. Foster and is now managed by R. Condit, S. Lao, and R. Perez under the Center for Tropical Forest Science and the Smithsonian Tropical Research Institute in Panama. Numerous organizations have provided funding, principally the U.S. National Science Foundation. Support for the establishment and monitoring of permanent plots of RLC and BF in Costa Rica was provided by grants from the Andrew W. Mellon Foundation, the US National Science Foundation (NSF DEB-0424767, NSF DEB-0639393 and NSF DEB-1147429), US NASA Terrestrial Ecology Program, and the University of Connecticut Research Foundation.

## AUTHOR CONTRIBUTIONS

The idea for this study was conceived during a workshop attended by LA, FB, RLC, DC, CEF, BH, CCJ, SK, JAM, RM, NN, LP, NR, MES, MvB and MTvdS. RLC, DC, DHD, DD, SJD, JMD, BF, JSH, JLHS, ORL, JAM, RM, and MvB contributed data. MES, NR and SK prepared forest inventory data for analysis and calculated demographic rates. MES analysed the data. MES wrote the first draft of the manuscript with support from NR. All authors contributed critically to the drafts and gave final approval for publication.

## DATA AVAILABILITY STATEMENT

Should the manuscript be accepted, the data supporting the results will be archived in an appropriate public repository (e.g. Dryad) and the data DOI will be included at the end of the article.

