## Supplementary_material for "Tree demographic strategies largely overlap across succession in Neotropical wet and dry forest communities"

### SUPPORTING INFORMATION

Table of Contents

1) Data Handling
   a) Sarapiquí
   b) Tirimbina
   c) Carbono
   d) Agua Salud
   e) BCNM
   f) BCI
   g) Yucatán Permanent plots
   h) Yucatán Conglomerates
   i) Oaxaca
2) Canopy layer assignment

Tables S1-S3

Figures S1-S17

References

#### 1. Data Handling

##### a) Sarapiquí

We use data from 7 secondary forest (SF) and 2 old growth forest (OGF) plots, each 1 ha in size (50 x 200 m). The minimum dbh is 5 cm. In 4 SF and 2 OGF plots, there is an additional census interval with sapling data (1-5 cm) from 0.5 ha (5 transects of 5 x 200 m per plot; Lindero Sur: 0.25 ha, 5 transects of 5 x 100 m). Lindero Sur new plot was not considered. We removed 22 duplicate entries in the 2017 census. We used last noted species names in case they changed or were missing. We removed lianas. In Cuatro Ríos, we corrected one x-coordinate (deemed 50 instead of 80). In Lindero Sur, y-coordinates exceeding 100 were subtracted 100. We discarded 42 remnant trees (22 in Finca el Bejuco, 9 in Lindero Sur, 6 in Tirimbina, 3 in Lindero el Peje secundario, 2 in Cuatro Ríos).

##### b) Tirimbina

We use data from four SF plots (Manú & Aceituno 1.16 ha, Bottarrama 1.6 ha, Arrozal 1987-2003 0.3 ha, 2006-2017 1.44 ha). Minimum dbh is generally 5 cm, but varies (see Table S1). We corrected species affiliation for three entries (1-63-35: Chrysoplyllum brenesii, 4-34-38: Beilschmiedia costaricensis, 4‑53‑35 Cestrum microcalyx) and treeID for one entry (subplot 74 in Arrozal does not exist; correct treeID for 4-74-35-1 is 4-73-35-1).

##### c) Carbono

We use data from 12 OGF plots each 0.5 ha in size. We only use plots from residual soils (plots L (flat) and P (slopes)) to be consistent with the Sarapiquí and Tirimbina secondary forest data. Minimum dbh is 10 cm. For one census interval, there is additional information on sapling data (1-10 cm) from one subplot per plot (0.01 ha each). We excluded trees with negative coordinates.

##### d) Agua Salud

We use data from 104 plots (referred to as transects) each 0.1 ha in size (20 x 50 m). Transects 1-4 were not considered due to differences in sampling design. Minimum dbh is 5 cm for one half of each transect and 1 cm for the other half. For 10 transects (73, 74, 79, 80, 91, 92, 97-100) exact age since abandonment could only be estimated to be between 50 and 80 years. We used an age of 50 years in 2009 for our analyses.

##### e) BCNM

We use data from 8 SF plots each 0.32 ha in size except Saino (0.18 ha). Minimum dbh is 5 cm. We removed 3 duplicate entries. We corrected census year = 2102 to 2002.

##### f) BCI

We use data from one 50 ha OGF plot. Minimum dbh is 1 cm. Trees with no dbh measurement but status “alive” were not considered. We used the mean census date of the census year in case the exact date of measurement was missing. We discarded trees with unknown location.

##### g) Yucatán Permanent plots

We use data from 9 SF plots each 0.1 ha (20 x 50 m) in size. Minimum dbh is generally 5 cm and 1 cm for ten 30 m² subplots per plot.

#### h) Yucatán Conglomerates

We use data from 20 SF plots, referred to as conglomerates. Each conglomerate consists of 4 circular 400 m² plots (11.28 m in radius) within a circular area of 1 ha. Minimum dbh is generally 7.5 cm and 2.5 cm for one 80 m² concentric subplot per plot. We corrected typing errors in conglomerate, plot, individual or stem numbers in 526 entries. We corrected a typing error in dbh measurement in 1 entry. We excluded 4 conglomerates and 7 subplots due to exceeding numbers of typing errors.

##### i) Oaxaca

We use data from 17 SF and 8 OGF plots each 0.04 ha (20 x 20 m) or 0.05 ha (20 x 25 m) in size (see Table S1). Minimum dbh is 5 cm for 25% (OGF) or 50% (SF) of each plot, 2.5cm for additional 50% (OGF) or 25% (SF) of each plot and 1 cm for the rest of each plot. OGF plots that were established on soils derived from limestone were not considered in order to be consistent with SF plots. We excluded lianas. Trees outside of plot borders were discarded. Trees noted dead but alive again in later censuses were omitted from those censuses until again noted alive.

#### 2. Canopy layer assignment

Trees were assigned to one of three canopy layers following the approach of Purves *et al.* (2008) and Bohlman & Pacala (2012). To do this, plots were divided into subplots of different size (see Table S1). Within each subplot, trees were sorted by size and assigned to the top canopy layer (layer 1) until the cumulative area of their crowns exceeded the subplot area. For the smallest tree to enter layer 1 (i.e. the top canopy layer), we checked whether >50% of its crown area would ‘fit’ into layer 1. If this was the case, we assigned it to layer 1, otherwise, we assigned it to layer 2. Smaller trees were assigned to layer 2 until the cumulative area of their crowns exceeded the subplot area, and so on. For each tree, we calculated its crown area from common (i.e. the same for all species) site-specific allometries.

For Costa Rica, the dbh – crown area allometric relationship was calculated based on measurements from 574 trees located in the Lindero el Peje secundario plot from the Sarapiquí dataset (Eqn 1).

(Equation 1)

$$crown area \left( m^{2} \right)=0.048*{dbh \left( mm \right)}^{1.205}$$

For Panama, the allometric equation from Bohlman & Pacala (2012) was used (Eqn 2).

(Equation 2)

$$crown area \left( m^{2} \right)=0.036*{dbh \left( mm \right)}^{1.281}$$

For Oaxaca, 29.662 crown area and dbh observations from 3.828 individuals were used to calculate allometric equations. Separate allometries were derived for trees (28.587 obs. from 3.609 individuals, Eqn 3) and cacti (1.075 from 219 individuals, Eqn 4).

(Equation 3)

$$crown area \left( m^{2} \right)=188.630*{ba \left( m^{2} \right)}^{0.578}$$

(Equation 4)

$$crown area \left( m^{2} \right)=29.633*{ba \left( m^{2} \right)}^{0.858}$$

For Yucatán, the tree allometry from Oaxaca was used (Eqn 3).

**Table S1**: All plots and census intervals used along with information on key references, previous land use, stand age, minimum dbh and areas.

| **Site** | **Dataset** | **Key references** | **Plot** | **Previous land use** | **Census interval** | **Stand age (years)** | **Minimum dbh (cm)** | **Area (ha)** | **Minimum dbh subplots (cm)** | **Area subplots (ha)** | **Subplot area for canopy layer assignment (m²)** |
| --- | --- | --- | --- | --- | --- | --- | --- | --- | --- | --- | --- |
| Costa Rica | Tirimbina | Chazdon *et al.* (2007)  Chazdon *et al.* (2010) | Arrozal | Rice cultivation  (1 year) | 1987-1992 | 1-6 | 5 | 0.30 |  |  | 100 |
|  |  |  |  |  | 1992-1998 | 6-12 | 5 | 0.30 |  |  | 100 |
|  |  |  |  |  | 1998-2003 | 12-17 | 5 | 0.30 |  |  | 100 |
|  |  |  |  |  | 2008-2013 | 22-27 | 5 | 1.44 |  |  | 800 |
|  |  |  |  |  | 2013-2017 | 27-31 | 5 | 1.44 |  |  | 800 |
|  |  |  | Aceituno | Cleared but not used | 1990-1995 | 18-23 | 5 | 1.16 |  |  | 800 |
|  |  |  |  |  | 1998-2003 | 26-31 | 5 | 1.16 |  |  | 800 |
|  |  |  |  |  | 2008-2013 | 36-41 | 5 | 1.16 |  |  | 800 |
|  |  |  |  |  | 2013-2017 | 41-45 | 5 | 1.16 |  |  | 800 |
|  |  |  | Botarrama | Cleared but not used | 1990-1995 | 28-33 | 10 | 1.60 |  |  | 800 |
|  |  |  |  |  | 1998-2003 | 36-41 | 10 | 1.60 |  |  | 800 |
|  |  |  |  |  | 2003-2008 | 41-46 | 10 | 1.60 |  |  | 800 |
|  |  |  |  |  | 2008-2013 | 46-51 | 5 | 1.60 |  |  | 800 |
|  |  |  |  |  | 2013-2017 | 51-55 | 5 | 1.60 |  |  | 800 |
|  |  |  | Manú | Cleared but not used | 1990-1995 | 30-35 | 10 | 1.16 |  |  | 800 |
|  |  |  |  |  | 1998-2003 | 38-43 | 10 | 1.16 |  |  | 800 |
|  |  |  |  |  | 2003-2008 | 43-48 | 10 | 1.16 |  |  | 800 |
|  |  |  |  |  | 2008-2013 | 48-53 | 5 | 1.16 |  |  | 800 |
|  |  |  |  |  | 2013-2017 | 53-57 | 5 | 1.16 |  |  | 800 |
|  | Sarapiquí | Letcher & Chazdon (2009) Chazdon *et al.* (2011) | Finca el Bejuco | Pasture | 2006-2011 | 12-17 | 5 | 1.00 | 1 | 0.50 | 625 |
|  |  |  |  |  | 2011-2016 | 17-22 | 5 | 1.00 |  |  | 625 |
|  |  |  | Juan Enriquez | Pasture | 2006-2011 | 12-17 | 5 | 1.00 | 1 | 0.50 | 625 |
|  |  |  |  |  | 2011-2016 | 17-22 | 5 | 1.00 |  |  | 625 |
|  |  |  | Lindero Sur | Pasture | 1997-2002 | 12-17 | 5 | 1.00 |  |  | 625 |
|  |  |  |  |  | 2002-2007 | 17-22 | 5 | 1.00 |  |  | 625 |
|  |  |  |  |  | 2007-2012 | 22-27 | 5 | 1.00 | 1 | 0.25 | 625 |
|  |  |  | Tirimbina | Pasture | 1997-2002 | 15-20 | 5 | 1.00 |  |  | 625 |
|  |  |  |  |  | 2002-2007 | 20-25 | 5 | 1.00 |  |  | 625 |
|  |  |  |  |  | 2007-2012 | 25-30 | 5 | 1.00 |  |  | 625 |
|  |  |  |  |  | 2012-2017 | 30-35 | 5 | 1.00 |  |  | 625 |
|  |  |  | Lindero el Peje Secundario | Pasture | 1997-2002 | 20-25 | 5 | 1.00 |  |  | 625 |
|  |  |  |  |  | 2002-2007 | 25-30 | 5 | 1.00 |  |  | 625 |
|  |  |  |  |  | 2007-2012 | 30-35 | 5 | 1.00 | 1 | 0.50 | 625 |
|  |  |  |  |  | 2012-2017 | 35-40 | 5 | 1.00 |  |  | 625 |
| **Site** | **Dataset** | **Key references** | **Plot** | **Previous land use** | **Census interval** | **Stand age (years)** | **Minimum dbh (cm)** | **Area (ha)** | **Minimum dbh subplots (cm)** | **Area subplots (ha)** | **Subplot area for canopy layer assignment (m²)** |
| Costa Rica | Sarapiquí |  | Cuatro Ríos | Pasture | 1997-2002 | 25-30 | 5 | 1.00 |  |  | 625 |
|  |  |  |  |  | 2002-2007 | 30-35 | 5 | 1.00 |  |  | 625 |
|  |  |  |  |  | 2007-2012 | 35-40 | 5 | 1.00 |  |  | 625 |
|  |  |  |  |  | 2012-2017 | 40-45 | 5 | 1.00 |  |  | 625 |
|  |  |  | Lindero el Peje Primario | - | 2007-2012 | OGF | 5 | 1.00 | 1 | 0.50 | 625 |
|  |  |  |  |  | 2012-2017 | OGF | 5 | 1.00 |  |  | 625 |
|  |  |  | Selva Verde | - | 2007-2012 | OGF | 5 | 1.00 | 1 | 0.50 | 625 |
|  |  |  |  |  | 2012-2017 | OGF | 5 | 1.00 |  |  | 625 |
|  | Carbono | Clark & Clark (2000) | L1-L6, P1-P6 (12 plots total) | - | 1997-2002 | OGF | 10 | 0.50 (each) |  |  | 625 |
|  |  |  |  |  | 2002-2007 | OGF | 10 | 0.50 (each) |  |  | 625 |
|  |  |  |  |  | 2007-2012 | OGF | 10 | 0.50 (each) | 1 | 0.01 (each) | 625 |
|  |  |  |  |  | 2012-2016 | OGF | 10 | 0.50 (each) |  |  | 625 |
| Panama | Agua Salud | van Breugel *et al.* (2013) Lai *et al.* (2017) | Transects 5-104, 131-134 | Pasture, shifting cultivation | 2009-2014 | 1-55 | 5 | 0.10 (each) | 1 | 0.05 (each) | 1000 |
|  | BCNM | Denslow & Guzman (2000) | Saino | Pasture, swidden agriculture | 2001-2011 | 30-40 | 5 | 0.18 |  |  | 800 |
|  |  |  | Pedro Gomez | Pasture, swidden agriculture | 2001-2011 | 30-40 | 5 | 0.32 |  |  | 800 |
|  |  |  | Enders | Plantation | 2001-2011 | 50-60 | 5 | 0.32 |  |  | 800 |
|  |  |  | Foster | Pasture, swidden agriculture | 2001-2011 | 50-60 | 5 | 0.32 |  |  | 800 |
|  |  |  | Poacher’s Point | Pasture, swidden agriculture | 2001-2011 | 80-90 | 5 | 0.32 |  |  | 800 |
|  |  |  | Bohio | Pasture | 2001-2011 | 80-90 | 5 | 0.32 |  |  | 800 |
|  |  |  | Barbour | Pasture | 2001-2011 | 110-120 | 5 | 0.32 |  |  | 800 |
|  |  |  | Pearson | Pasture | 2001-2011 | 110-120 | 5 | 0.32 |  |  | 800 |
|  | BCI | Hubbell & Foster (1983)  Condit (1998) Hubbell *et al.* (1999) | BCI | - | 1985-1990 | OGF | 1 | 50.00 |  |  | 976.5625 |
|  |  |  |  |  | 1990-1995 | OGF | 1 | 50.00 |  |  | 976.5625 |
|  |  |  |  |  | 1995-2000 | OGF | 1 | 50.00 |  |  | 976.5625 |

| **Site** | **Dataset** | **Key references** | **Plot** | **Previous land use** | **Census interval** | **Stand age (years)** | **Minimum dbh (cm)** | **Area (ha)** | **Minimum dbh subplots (cm)** | **Area subplots (ha)** | **Subplot area for canopy layer assignment (m²)** |
| --- | --- | --- | --- | --- | --- | --- | --- | --- | --- | --- | --- |
| Panama | BCI |  | BCI |  | 2000-2005 | OGF | 1 | 50.00 |  |  | 976.5625 |
|  |  |  |  |  | 2005-2010 | OGF | 1 | 50.00 |  |  | 976.5625 |
|  |  |  |  |  | 2010-2015 | OGF | 1 | 50.00 |  |  | 976.5625 |
| Yucatán | Permanent Plots | Saenz-Pedroza *et al.* (2020) | 3A | Slash-and-burn agriculture | 2009-2014 | 3-8 | 5 | 0.10 | 1 | 0.03 | 100 |
|  |  |  | 3B |  | 2009-2014 | 3-8 | 5 | 0.10 | 1 | 0.03 | 100 |
|  |  |  | 5C |  | 2009-2014 | 5-10 | 5 | 0.10 | 1 | 0.03 | 100 |
|  |  |  | 15B |  | 2005-2010 | 14-19 | 5 | 0.10 | 1 | 0.03 | 100 |
|  |  |  |  |  | 2010-2015 | 19-24 | 5 | 0.10 | 1 | 0.03 | 100 |
|  |  |  | 15C |  | 2005-2010 | 15-20 | 5 | 0.10 | 1 | 0.03 | 100 |
|  |  |  |  |  | 2010-2015 | 20-25 | 5 | 0.10 | 1 | 0.03 | 100 |
|  |  |  | 17A |  | 2005-2010 | 16-21 | 5 | 0.10 | 1 | 0.03 | 100 |
|  |  |  |  |  | 2010-2015 | 21-26 | 5 | 0.10 | 1 | 0.03 | 100 |
|  |  |  | 60B |  | 2005-2010 | 50-55 | 5 | 0.10 | 1 | 0.03 | 100 |
|  |  |  |  |  | 2010-2015 | 55-60 | 5 | 0.10 | 1 | 0.03 | 100 |
|  |  |  | 60A |  | 2005-2010 | 55-60 | 5 | 0.10 | 1 | 0.03 | 100 |
|  |  |  |  |  | 2010-2015 | 60-65 | 5 | 0.10 | 1 | 0.03 | 100 |
|  |  |  | 60C |  | 2005-2010 | 60-65 | 5 | 0.10 | 1 | 0.03 | 100 |
|  |  |  |  |  | 2010-2015 | 65-70 | 5 | 0.10 | 1 | 0.03 | 100 |
|  | Conglomerate Data | Hernández-Stefanoni *et al.* (2014) | 6-9, 11-17, 19, 25-32 | Slash-and-burn agriculture | 2013-2018 | 10-85 | 7.5 | 0.16 (each) | 2.5 | 0.032 | 400 |
| Oaxaca | Nizanda | Lebrija-Trejos *et al.* (2008)  Pérez-García *et al.* (2010)  Lebrija-Trejos *et al.* (2010)  Lebrija-Trejos *et al.* (2011) Muñoz *et al.* (2021) | MAR | Few years of cultivation without pasture use | 2004-2009 | 4-9 | 5 | 0.04 | 2.5 / 1 | 0.02 / 0.01 | 100 |
|  |  |  | TOB |  | 2006-2011 | 4-9 | 5 | 0.04 | 2.5 / 1 | 0.02 / 0.01 | 100 |
|  |  |  |  |  | 2011-2016 | 9-14 | 5 | 0.04 | 2.5 / 1 | 0.02 / 0.01 | 100 |
|  |  |  | TOA |  | 2008-2013 | 5-10 | 5 | 0.04 | 2.5 / 1 | 0.02 / 0.01 | 100 |
|  |  |  |  |  | 2013-2018 | 10-15 | 5 | 0.04 | 2.5 / 1 | 0.02 / 0.01 | 100 |
|  |  |  | HIL |  | 2003-2008 | 5-10 | 5 | 0.04 | 2.5 / 1 | 0.02 / 0.01 | 100 |
|  |  |  |  |  | 2008-2013 | 10-15 | 5 | 0.04 | 2.5 / 1 | 0.02 / 0.01 | 100 |
|  |  |  |  |  | 2013-2018 | 15-20 | 5 | 0.04 | 2.5 / 1 | 0.02 / 0.01 | 100 |
|  |  |  | DIA |  | 2003-2008 | 7-12 | 5 | 0.04 | 2.5 / 1 | 0.02 / 0.01 | 100 |
|  |  |  |  |  | 2008-2013 | 12-17 | 5 | 0.04 | 2.5 / 1 | 0.02 / 0.01 | 100 |
|  |  |  |  |  | 2013-2018 | 17-22 | 5 | 0.04 | 2.5 / 1 | 0.02 / 0.01 | 100 |

| **Site** | **Dataset** | **Key references** | **Plot** | **Previous land use** | **Census interval** | **Stand age (years)** | **Minimum dbh (cm)** | **Area (ha)** | **Minimum dbh subplots (cm)** | **Area subplots (ha)** | **Subplot area for canopy layer assignment (m²)** |
| --- | --- | --- | --- | --- | --- | --- | --- | --- | --- | --- | --- |
| Oaxaca |  |  | FID | Few years of cultivation without pasture use | 2003-2008 | 9-14 | 5 | 0.04 | 2.5 / 1 | 0.02 / 0.01 | 100 |
|  |  |  |  |  | 2008-2013 | 14-19 | 5 | 0.04 | 2.5 / 1 | 0.02 / 0.01 | 100 |
|  |  |  |  |  | 2013-2018 | 19-24 | 5 | 0.04 | 2.5 / 1 | 0.02 / 0.01 | 100 |
|  |  |  | ABE |  | 2003-2008 | 10-15 | 5 | 0.04 | 2.5 / 1 | 0.02 / 0.01 | 100 |
|  |  |  |  |  | 2008-2013 | 15-20 | 5 | 0.04 | 2.5 / 1 | 0.02 / 0.01 | 100 |
|  |  |  |  |  | 2013-2018 | 20-25 | 5 | 0.04 | 2.5 / 1 | 0.02 / 0.01 | 100 |
|  |  |  | BES |  | 2003-2008 | 11-16 | 5 | 0.04 | 2.5 / 1 | 0.02 / 0.01 | 100 |
|  |  |  | ESS |  | 2003-2008 | 16-21 | 5 | 0.04 | 2.5 / 1 | 0.02 / 0.01 | 100 |
|  |  |  |  |  | 2008-2013 | 21-26 | 5 | 0.04 | 2.5 / 1 | 0.02 / 0.01 | 100 |
|  |  |  |  |  | 2013-2018 | 26-31 | 5 | 0.04 | 2.5 / 1 | 0.02 / 0.01 | 100 |
|  |  |  | ISC |  | 2004-2009 | 19-24 | 5 | 0.04 | 2.5 / 1 | 0.02 / 0.01 | 100 |
|  |  |  |  |  | 2009-2014 | 24-29 | 5 | 0.04 | 2.5 / 1 | 0.02 / 0.01 | 100 |
|  |  |  |  |  | 2014-2018 | 29-33 | 5 | 0.04 | 2.5 / 1 | 0.02 / 0.01 | 100 |
|  |  |  | ISP |  | 2003-2008 | 23-28 | 5 | 0.04 | 2.5 / 1 | 0.02 / 0.01 | 100 |
|  |  |  |  |  | 2009-2014 | 29-34 | 5 | 0.04 | 2.5 / 1 | 0.02 / 0.01 | 100 |
|  |  |  |  |  | 2014-2018 | 34-38 | 5 | 0.04 | 2.5 / 1 | 0.02 / 0.01 | 100 |
|  |  |  | RIC |  | 2003-2008 | 30-35 | 5 | 0.04 | 2.5 / 1 | 0.02 / 0.01 | 100 |
|  |  |  |  |  | 2008-2013 | 35-40 | 5 | 0.04 | 2.5 / 1 | 0.02 / 0.01 | 100 |
|  |  |  |  |  | 2013-2018 | 40-45 | 5 | 0.04 | 2.5 / 1 | 0.02 / 0.01 | 100 |
|  |  |  | MAL |  | 2003-2008 | 36-41 | 5 | 0.04 | 2.5 / 1 | 0.02 / 0.01 | 100 |
|  |  |  |  |  | 2008-2013 | 41-46 | 5 | 0.04 | 2.5 / 1 | 0.02 / 0.01 | 100 |
|  |  |  |  |  | 2013-2018 | 46-51 | 5 | 0.04 | 2.5 / 1 | 0.02 / 0.01 | 100 |
|  |  |  | SEP |  | 2003-2008 | 40-45 | 5 | 0.04 | 2.5 / 1 | 0.02 / 0.01 | 100 |
|  |  |  |  |  | 2008-2013 | 45-50 | 5 | 0.04 | 2.5 / 1 | 0.02 / 0.01 | 100 |
|  |  |  |  |  | 2013-2018 | 50-55 | 5 | 0.04 | 2.5 / 1 | 0.02 / 0.01 | 100 |
|  |  |  | DIV |  | 2005-2010 | 56-61 | 5 | 0.04 | 2.5 / 1 | 0.02 / 0.01 | 100 |
|  |  |  | JUL |  | 2005-2010 | 60-65 | 5 | 0.04 | 2.5 / 1 | 0.02 / 0.01 | 100 |
|  |  |  |  |  | 2010-2015 | 65-70 | 5 | 0.04 | 2.5 / 1 | 0.02 / 0.01 | 100 |
|  |  |  | LE1 | - | 2008-2013 | OGF | 5 | 0.05 | 2.5 / 1 | 0.0375 / 0.0125 | 125 |
|  |  |  |  |  | 2013-2018 | OGF | 5 | 0.05 | 2.5 / 1 | 0.0375 / 0.0125 | 125 |

| **Site** | **Dataset** | **Key references** | **Plot** | **Previous land use** | **Census interval** | **Stand age (years)** | **Minimum dbh (cm)** | **Area (ha)** | **Minimum dbh subplots (cm)** | **Area subplots (ha)** | **Subplot area for canopy layer assignment (m²)** |
| --- | --- | --- | --- | --- | --- | --- | --- | --- | --- | --- | --- |
| Oaxaca |  |  | LE2 | Few years of cultivation without pasture use | 2008-2013 | OGF | 5 | 0.05 | 2.5 / 1 | 0.0375 / 0.0125 | 125 |
|  |  |  |  |  | 2013-2018 | OGF | 5 | 0.05 | 2.5 / 1 | 0.0375 / 0.0125 | 125 |
|  |  |  | LPA |  | 2008-2013 | OGF | 5 | 0.05 | 2.5 / 1 | 0.0375 / 0.0125 | 125 |
|  |  |  |  |  | 2013-2018 | OGF | 5 | 0.05 | 2.5 / 1 | 0.0375 / 0.0125 | 125 |
|  |  |  | ISA |  | 2008-2013 | OGF | 5 | 0.05 | 2.5 / 1 | 0.0375 / 0.0125 | 125 |
|  |  |  |  |  | 2013-2018 | OGF | 5 | 0.05 | 2.5 / 1 | 0.0375 / 0.0125 | 125 |
|  |  |  | SEL |  | 2003-2008 | OGF | 5 | 0.04 | 2.5 / 1 | 0.02 / 0.01 | 100 |
|  |  |  |  |  | 2008-2013 | OGF | 5 | 0.04 | 2.5 / 1 | 0.02 / 0.01 | 100 |
|  |  |  |  |  | 2013-2018 | OGF | 5 | 0.04 | 2.5 / 1 | 0.02 / 0.01 | 100 |
|  |  |  | TEM |  | 2008-2013 | OGF | 5 | 0.05 | 2.5 / 1 | 0.0375 / 0.0125 | 125 |
|  |  |  |  |  | 2013-2018 | OGF | 5 | 0.05 | 2.5 / 1 | 0.0375 / 0.0125 | 125 |
|  |  |  | ZA1 |  | 2008-2013 | OGF | 5 | 0.05 | 2.5 / 1 | 0.0375 / 0.0125 | 125 |
|  |  |  |  |  | 2013-2018 | OGF | 5 | 0.05 | 2.5 / 1 | 0.0375 / 0.0125 | 125 |
|  |  |  | ZA2 |  | 2008-2013 | OGF | 5 | 0.05 | 2.5 / 1 | 0.0375 / 0.0125 | 125 |
|  |  |  |  |  | 2013-2018 | OGF | 5 | 0.05 | 2.5 / 1 | 0.0375 / 0.0125 | 125 |

**Table S2**: Number of observations, individuals and species that we calculated demographic rates for in early successional forests (ESF), late successional forests (LSF) and old-growth forests (OGF) for the four sites.

|  | **ESF** | | | | | **LSF** | | | | | **OGF** | | | | |
| --- | --- | --- | --- | --- | --- | --- | --- | --- | --- | --- | --- | --- | --- | --- | --- |
|  | **Nr. of observations** | | | **Nr. of indivi-duals** | **Nr. of species** | **Nr. of observations** | | | **Nr. of indivi-duals** | **Nr. of species** | **Nr. of observations** | | | **Nr. of indivi-duals** | **Nr. of species** |
|  | **Growth** | **Mortality** | **Recruit-ment** |  |  | **Growth** | **Mortality** | **Recruit-ment** |  |  | **Growth** | **Mortality** | **Recruit-ment** |  |  |
| **Costa Rica** | 17,387 | 22,526 | 1,053 | 14,662 | 348 | 16,411 | 18,876 | 86 | 7,093 | 334 | 11,975 | 13,578 | 236 | 5,636 | 329 |
| **Panama** | 18,783 | 28,968 | 9,186 | 39,812 | 324 | 5,158 | 6,264 | 479 | 7,375 | 334 | 1,102,739 | 1,278,550 | 153,821 | 265,141 | 304 |
| **Yucatán** | 4,628 | 5,596 | 248 | 5,447 | 131 | 3,737 | 4,234 | 101 | 3,775 | 123 | - | - | - |  | - |
| **Oaxaca** | 1,381 | 2,036 | 275 | 1,198 | 82 | 1,318 | 1,730 | 119 | 955 | 76 | 1,758 | 2,093 | 116 | 1,149 | 89 |

**Table S3**: Number of species included in hypervolume calculations for early successional forests (ESF), late successional forests (LSF) and old-growth forests (OGF) for the four sites for the hypervolume spanning growth and mortality.

|  | **ESF** | | | **LSF** | | | **OGF** | | |
| --- | --- | --- | --- | --- | --- | --- | --- | --- | --- |
|  | **Layer 1** | **Layer 2** | **Layer 3** | **Layer 1** | **Layer 2** | **Layer 3** | **Layer 1** | **Layer 2** | **Layer 3** |
| Costa Rica | 52 | 98 | 58 | 32 | 88 | 72 | 78 | 107 | 46 |
| Panama | 61 | 99 | 75 | 18 | 50 | 68 | 159 | 234 | 245 |
| Yucatán | 40 | 23 | 22 | 22 | 26 | 40 |  |  |  |
| Oaxaca | 5 | 14 | 7 | 9 | 19 | 23 | 5 | 23 | 28 |

**Table S4**: Number of species included in hypervolume calculations for early successional forests (ESF), late successional forests (LSF) and old-growth forests (OGF) for the four sites for the hypervolume spanning growth and recruitment.

|  | **ESF** | | | **LSF** | | | **OGF** | | |
| --- | --- | --- | --- | --- | --- | --- | --- | --- | --- |
|  | **Layer 1** | **Layer 2** | **Layer 3** | **Layer 1** | **Layer 2** | **Layer 3** | **Layer 1** | **Layer 2** | **Layer 3** |
| Costa Rica | 27 | 67 | 39 | 8 | 31 | 20 | 45 | 81 | 36 |
| Panama | 77 | 231 | 96 | 6 | 87 | 57 | 164 | 266 | 243 |
| Yucatán | 20 | 29 | 17 | 14 | 27 | 24 |  |  |  |
| Oaxaca | 8 | 45 | 9 | 6 | 29 | 22 | 11 | 29 | 25 |

**Table S5**: Number of species included in hypervolume calculations for early successional forests (ESF), late successional forests (LSF) and old-growth forests (OGF) for the four sites for the hypervolume spanning mortality and recruitment.

|  | **ESF** | | | **LSF** | | | **OGF** | | |
| --- | --- | --- | --- | --- | --- | --- | --- | --- | --- |
|  | **Layer 1** | **Layer 2** | **Layer 3** | **Layer 1** | **Layer 2** | **Layer 3** | **Layer 1** | **Layer 2** | **Layer 3** |
| Costa Rica | 25 | 44 | 31 | 7 | 16 | 14 | 38 | 47 | 33 |
| Panama | 56 | 94 | 75 | 5 | 23 | 48 | 157 | 229 | 241 |
| Yucatán | 19 | 15 | 14 | 9 | 12 | 21 |  |  |  |
| Oaxaca | 5 | 12 | 7 | 4 | 11 | 13 | 4 | 15 | 22 |

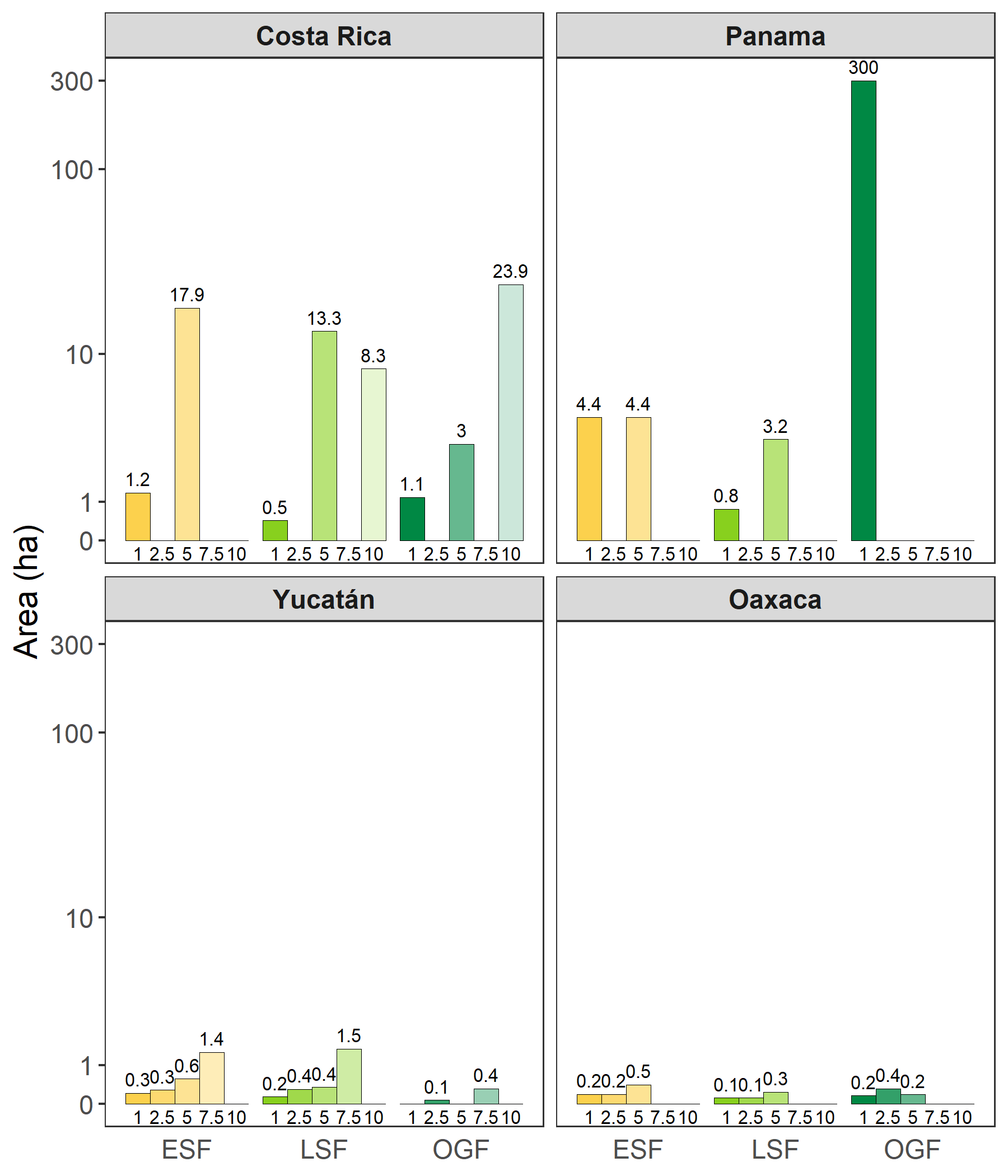

**Figure S1**: Sampling areas per minimum dbh threshold (cm) for early successional forests (ESF), late successional forests (LSF) and old-growth forests (OGF) for the four sites. Sampling area corresponds to the plot area * number of census intervals, e.g. OGF in Panama consists of 6 census intervals of the 50-ha plot at Barro Colorado Island, which has a minimum dbh threshold of 1 cm. In Yucatán, data from OGF was not available to sufficient extent and therefore not used subsequently.

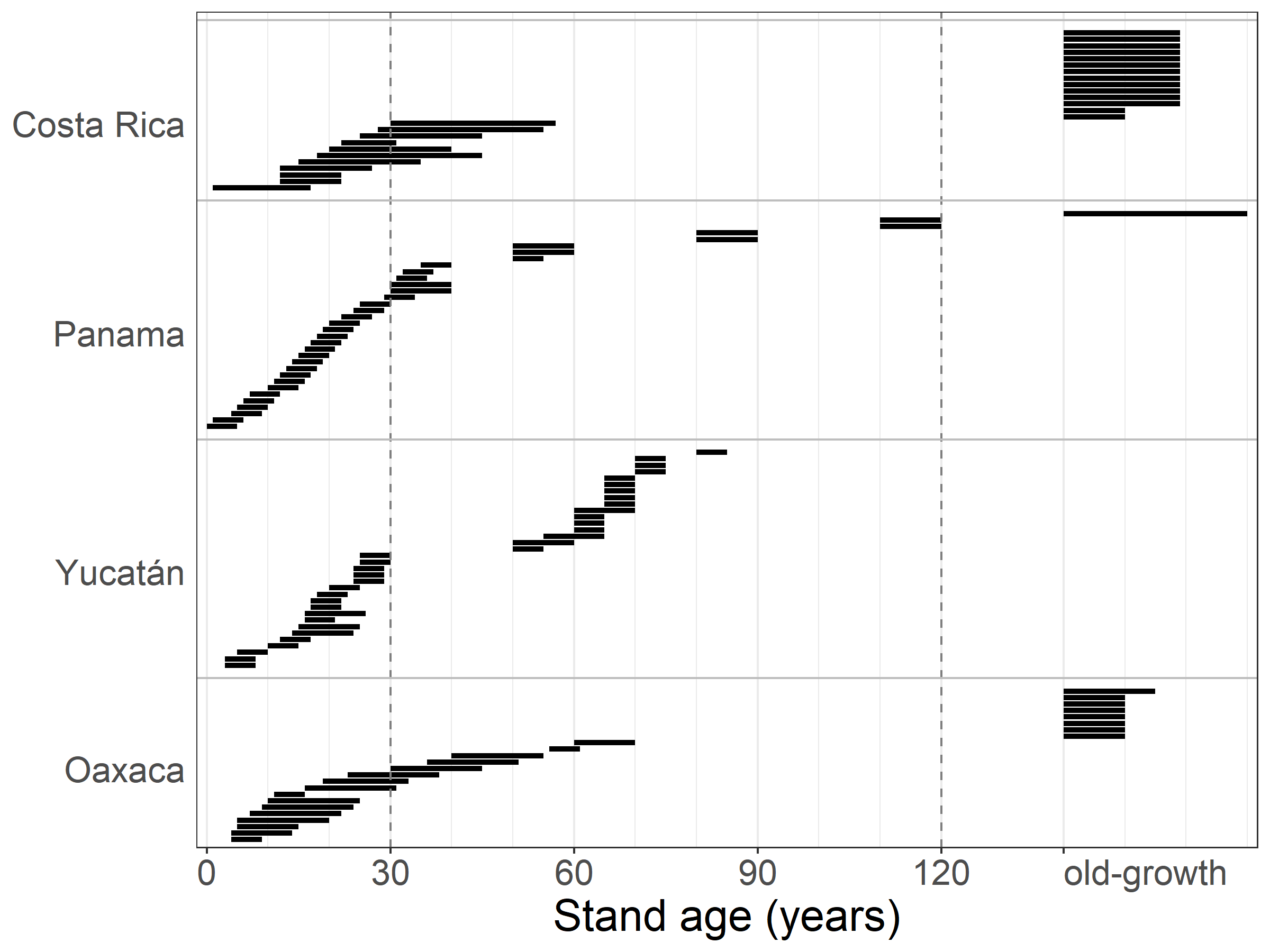

**Figure S2:** Stand age and census intervals of all plots included in this study. Each horizontal line corresponds to one or (in case there were too many) multiple plots in the chronosequence. Details are given in Table S1.

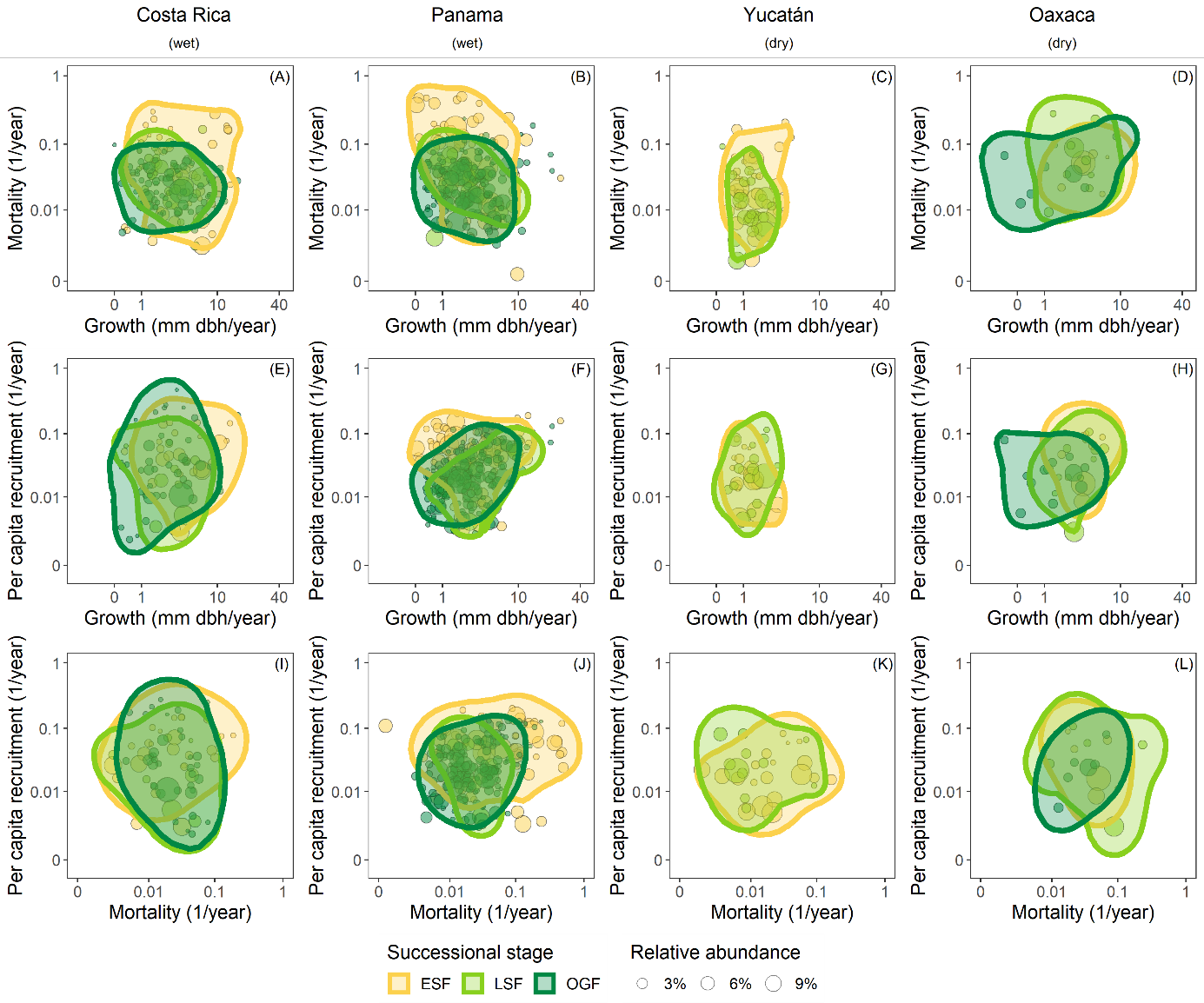

**Figure S3**: Demographic spaces for different pairs of demographic rates (A-D: growth-mortality, E-H: growth-recruitment, I-L: mortality-recruitment) for all sites and successional stages (ESF = early successional forest, LSF = late successional forest, OGF = old-growth forest) represented by two-dimensional hypervolumes. All axes are log-transformed (Ln). **Growth and mortality rates are from individuals assigned to canopy layer 1**. Hypervolume boundaries represent the smallest volume that captures a fraction of 80% of the total gaussian probability densities. All species contributed equally to the hypervolume calculation. Points represent species, point sizes indicate relative abundances within the successional stage.

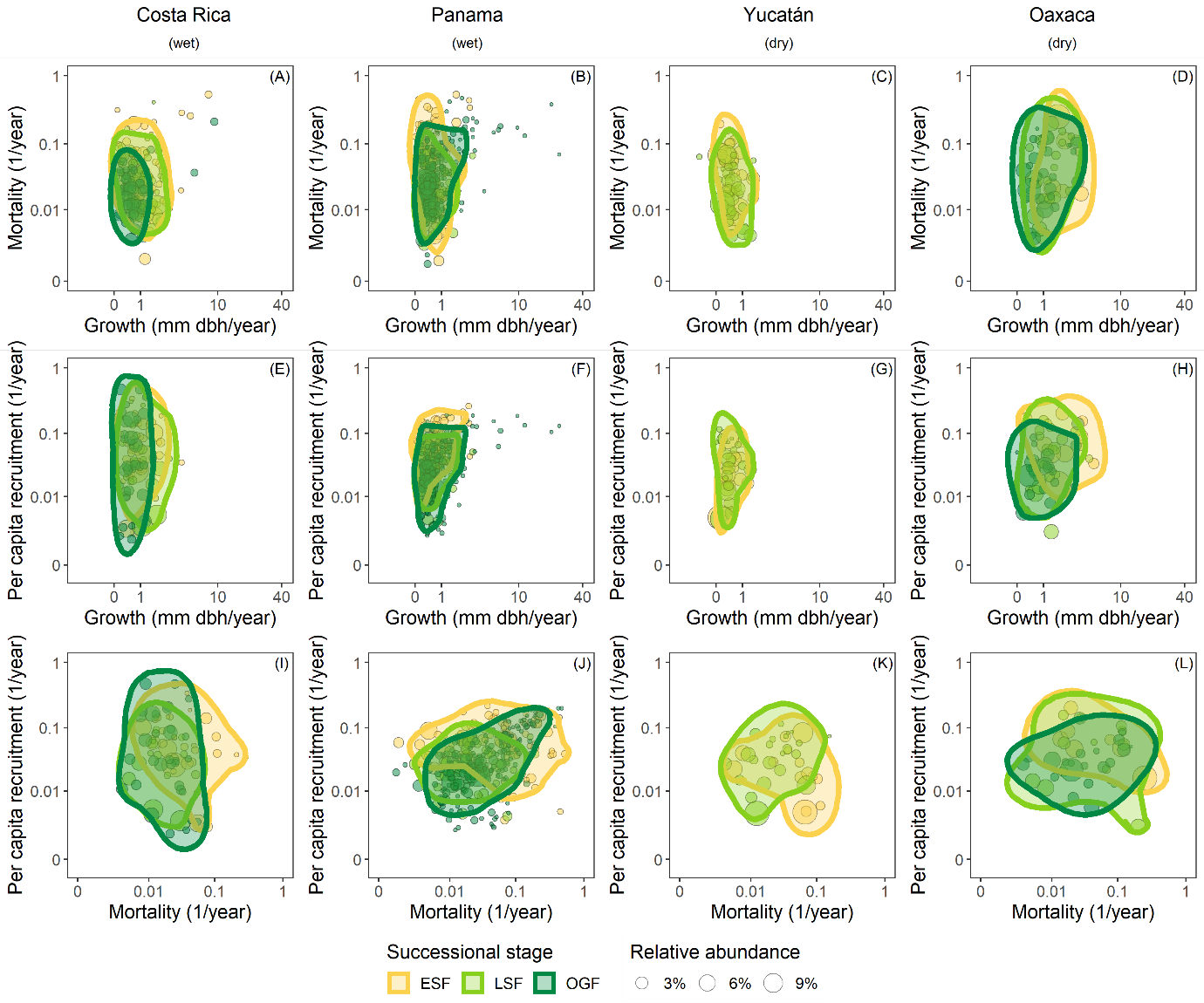

**Figure S4**: Demographic spaces for different pairs of demographic rates (A-D: growth-mortality, E-H: growth-recruitment, I-L: mortality-recruitment) for all sites and successional stages (ESF = early successional forest, LSF = late successional forest, OGF = old-growth forest) represented by two-dimensional hypervolumes. All axes are log-transformed (Ln). **Growth and mortality rates are from individuals assigned to canopy layer 3.** Hypervolume boundaries represent the smallest volume that captures a fraction of 80% of the total gaussian probability densities. All species contributed equally to the hypervolume calculation. Points represent species, point sizes indicate relative abundances within the successional stage.

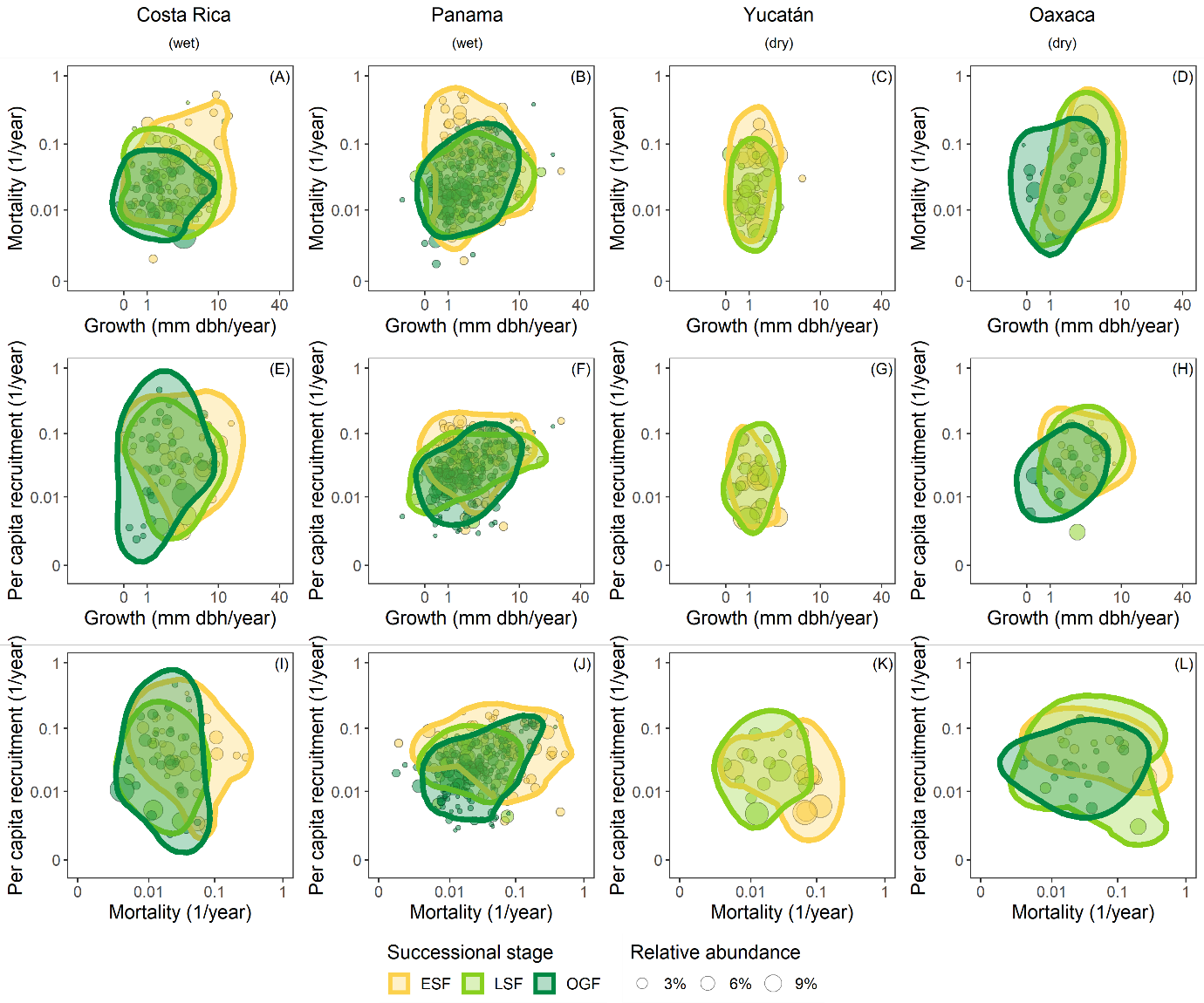

**Figure S5**: Demographic spaces for different pairs of demographic rates (A-D: growth-mortality, E-H: growth-recruitment, I-L: mortality-recruitment) for all sites and successional stages (ESF = early successional forest, LSF = late successional forest, OGF = old-growth forest) represented by two-dimensional hypervolumes. All axes are log-transformed (Ln). **Growth rates are from individuals assigned to canopy layer 1; mortality rates are from individuals assigned to canopy layer 3**. Hypervolume boundaries represent the smallest volume that captures a fraction of 80% of the total gaussian probability densities. All species contributed equally to the hypervolume calculation. Points represent species, point sizes indicate relative abundances within the successional stage.

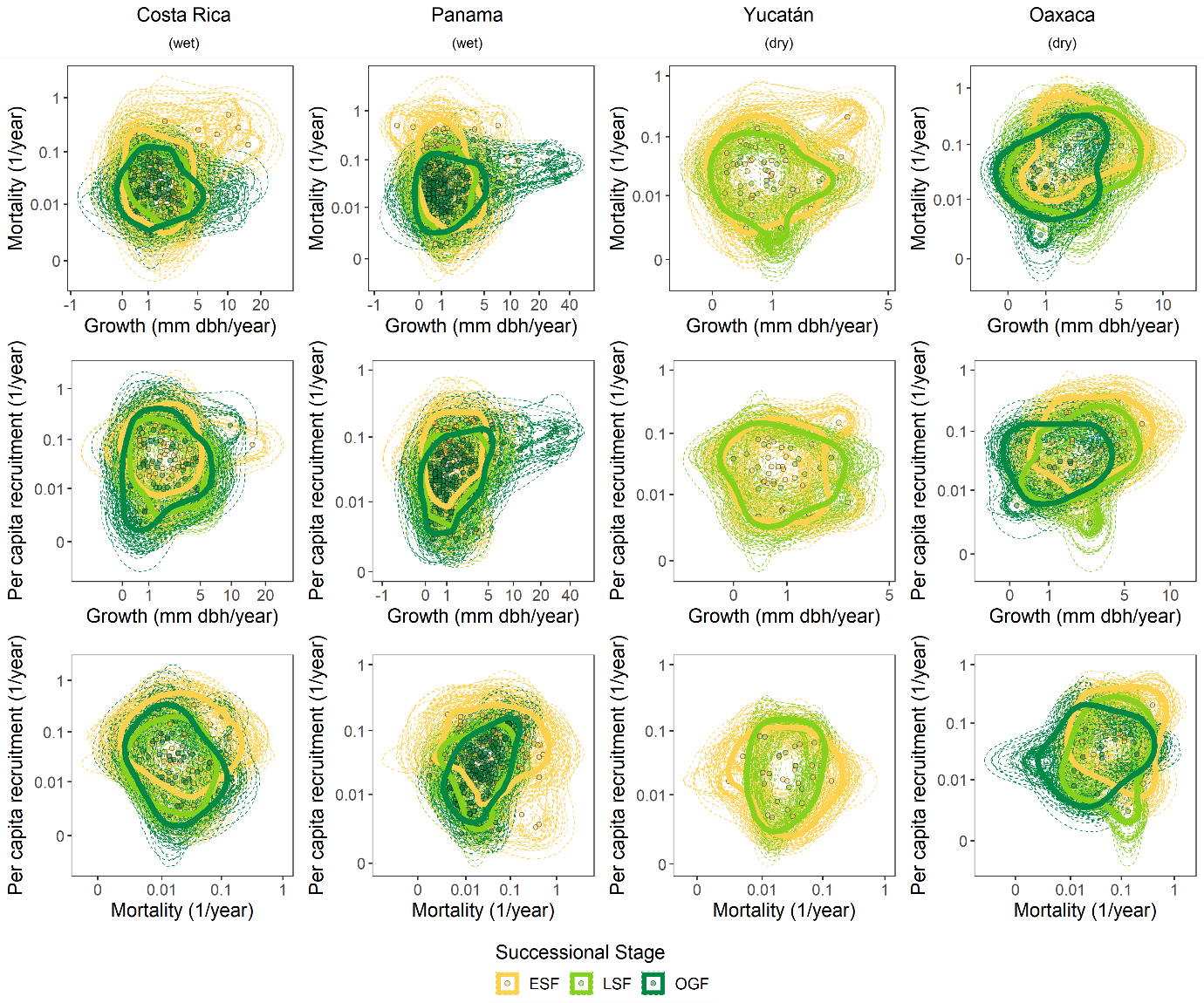

**Figure S6**: Original and rarefied demographic spaces for different pairs of demographic rates (A-D: growth-mortality, E-H: growth-recruitment, I-L: mortality-recruitment) for all sites and successional stages (ESF = early successional forest, LSF = late successional forest, OGF = old-growth forest) represented by two-dimensional hypervolumes. All axes are log-transformed (Ln). Growth and mortality rates are from individuals assigned to canopy layer 2. Hypervolume boundaries represent the smallest volume that captures a fraction of 80% of the total gaussian probability densities. All species contributed equally to the hypervolume calculation. Bold solid lines are the original demographic spaces; each dashed line is one of 100 bootstrap replicates with 10 randomly drawn species; points represent species.

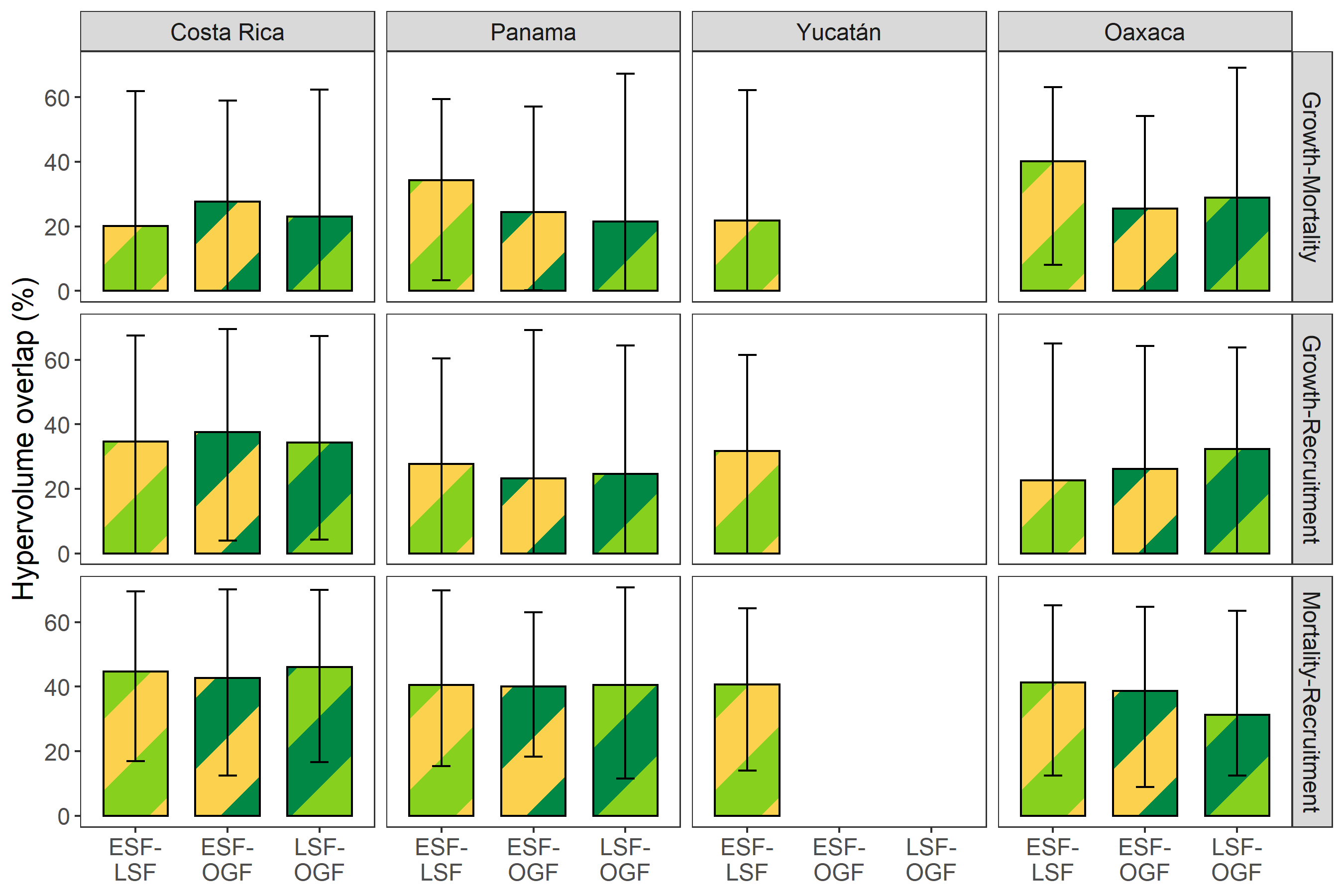

**Figure S7**: Overlap statistics of the two-dimensional hypervolumes (i.e. areas) representing the ranges of demographic strategies for different pairs of demographic rates and successional stages (ESF = early successional forest, LSF = late successional forest, OGF = old-growth forest). Colored bars represent the median rarefied and bootstrapped values, error bars represent 95% confidence intervals (r = 100 replicates, n = 10 species).

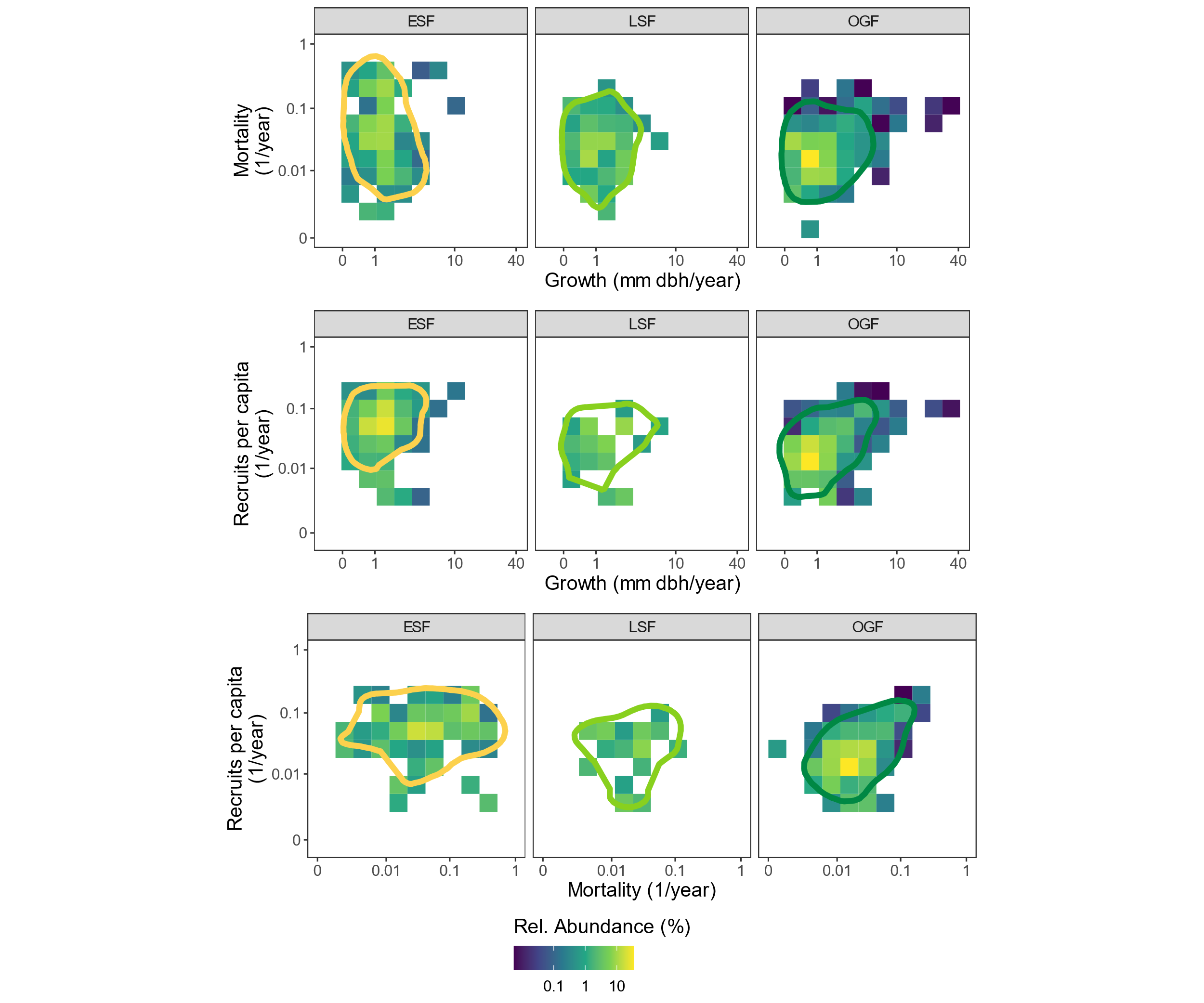

**Figure S8:** Demographic spaces for different pairs of demographic rates for **Costa Rica** separately for each successional stage represented by two-dimensional hypervolumes with underlying heatmaps depicting relative abundance in terms of number of trees per successional stage. All axes are log-transformed (Ln). Growth and mortality rates are from individuals assigned to canopy layer 2. Hypervolume boundaries represent the smallest volume that captures a fraction of 80% of the total gaussian probability densities. All species contributed equally to the hypervolume calculation.

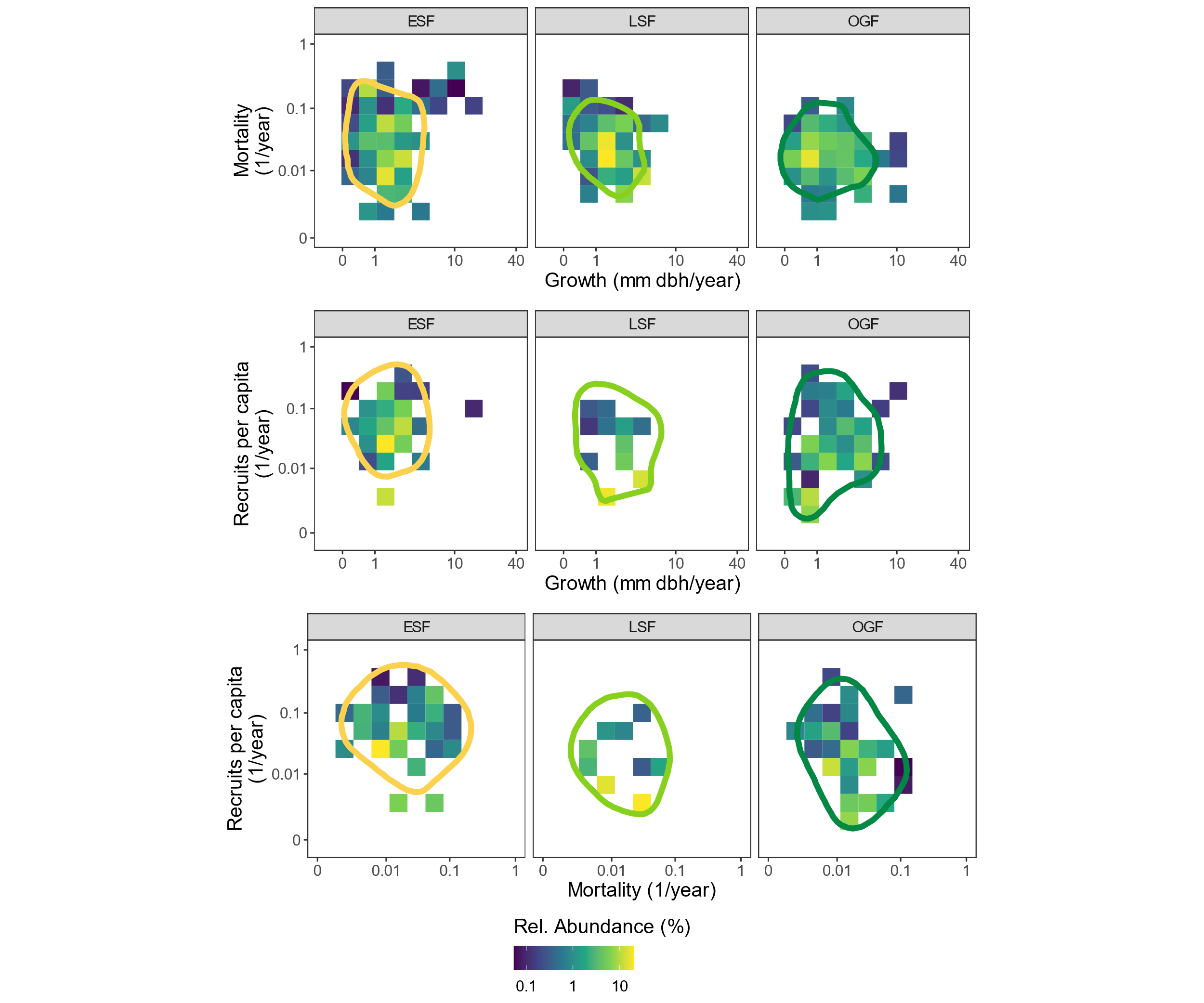

**Figure S9:** Demographic spaces for different pairs of demographic rates for **Panama** separately for each successional stage represented by two-dimensional hypervolumes with underlying heatmaps depicting relative abundance in terms of number of trees per successional stage. All axes are log-transformed (Ln). Growth and mortality rates are from individuals assigned to canopy layer 2. Hypervolume boundaries represent the smallest volume that captures a fraction of 80% of the total gaussian probability densities. All species contributed equally to the hypervolume calculation.

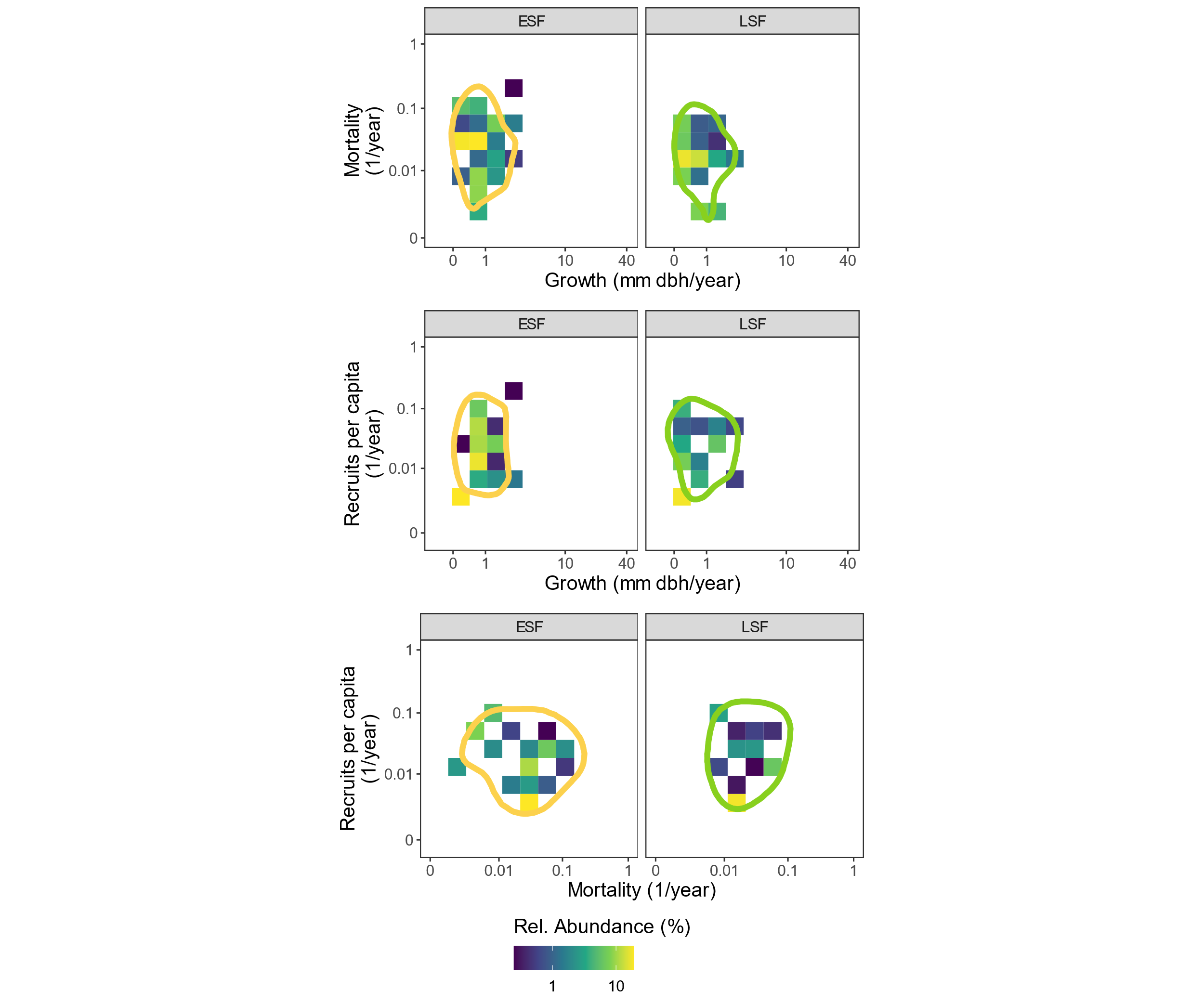

**Figure S10:** Demographic spaces for different pairs of demographic rates for **Yucatán** separately for each successional stage represented by two-dimensional hypervolumes with underlying heatmaps depicting relative abundance in terms of number of trees per successional stage. All axes are log-transformed (Ln). Growth and mortality rates are from individuals assigned to canopy layer 2. Hypervolume boundaries represent the smallest volume that captures a fraction of 80% of the total gaussian probability densities. All species contributed equally to the hypervolume calculation.

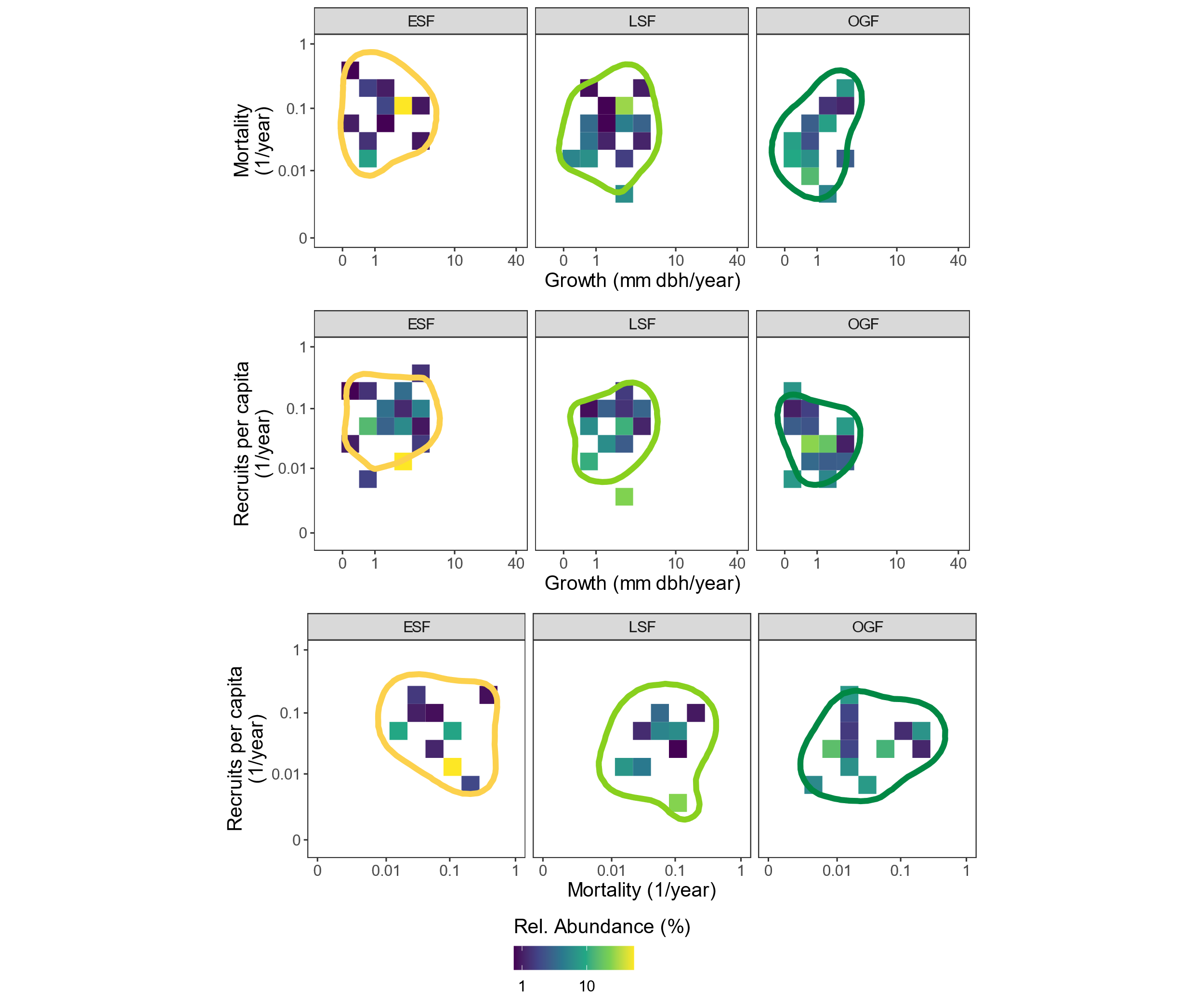

**Figure S11:** Demographic spaces for different pairs of demographic rates for **Oaxaca** separately for each successional stage represented by two-dimensional hypervolumes with underlying heatmaps depicting relative abundance in terms of number of trees per successional stage. All axes are log-transformed (Ln). Growth and mortality rates are from individuals assigned to canopy layer 2. Hypervolume boundaries represent the smallest volume that captures a fraction of 80% of the total gaussian probability densities. All species contributed equally to the hypervolume calculation.

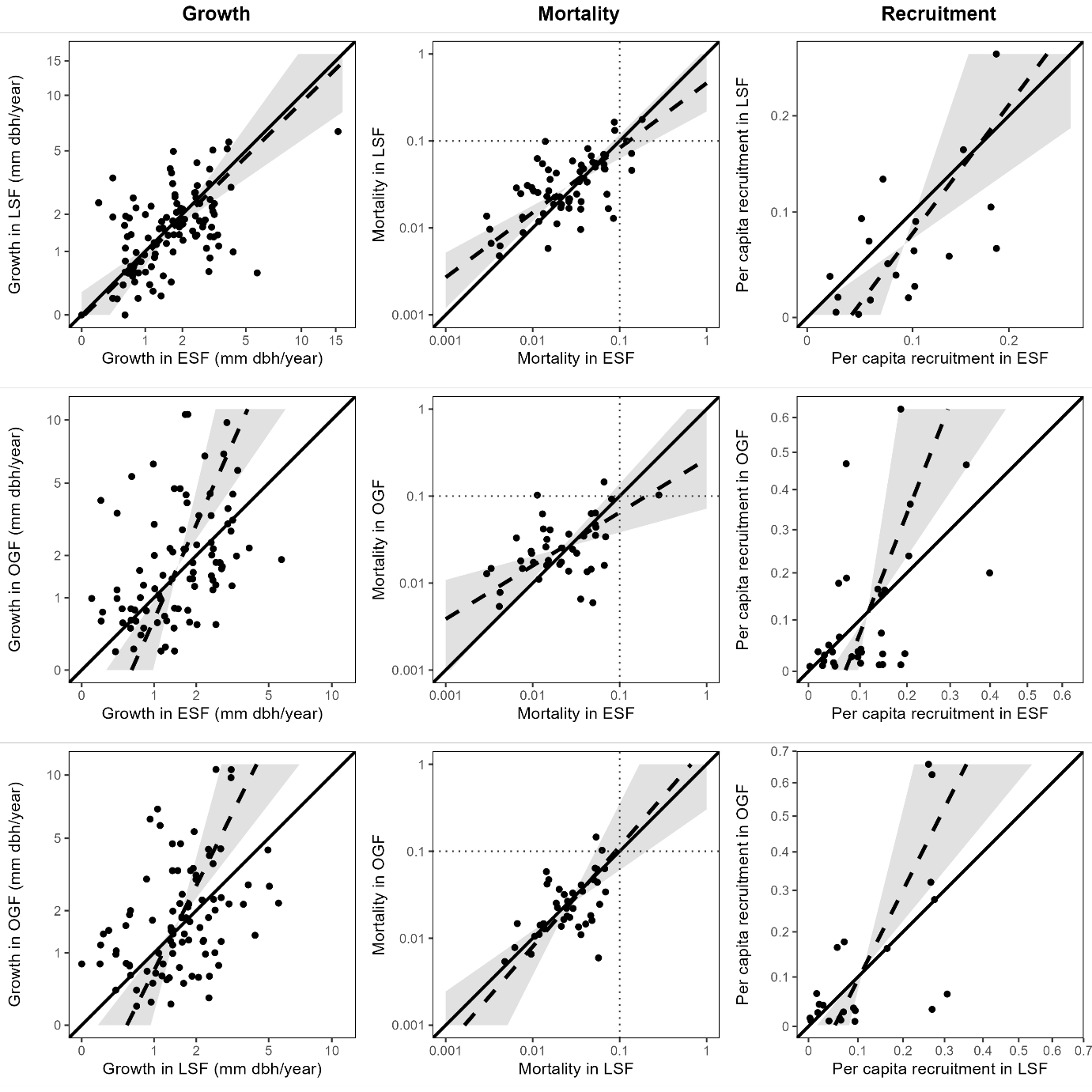

**Figure S12**: Major axis regressions for species’ demographic rates in early successional forests (ESF), late successional forests (LSF) and old-growth forests (OGF) in **Costa Rica**. Each point represents a species and its demographic rate in the respective successional stage. Solid lines represent the 1:1-line, dashed lines represent the regression lines and areas highlighted in grey represent the confidence intervals. Growth and mortality rates are from canopy layer 2. Only species with at least 5 observations for growth and survival in both successional stages were included, respectively.

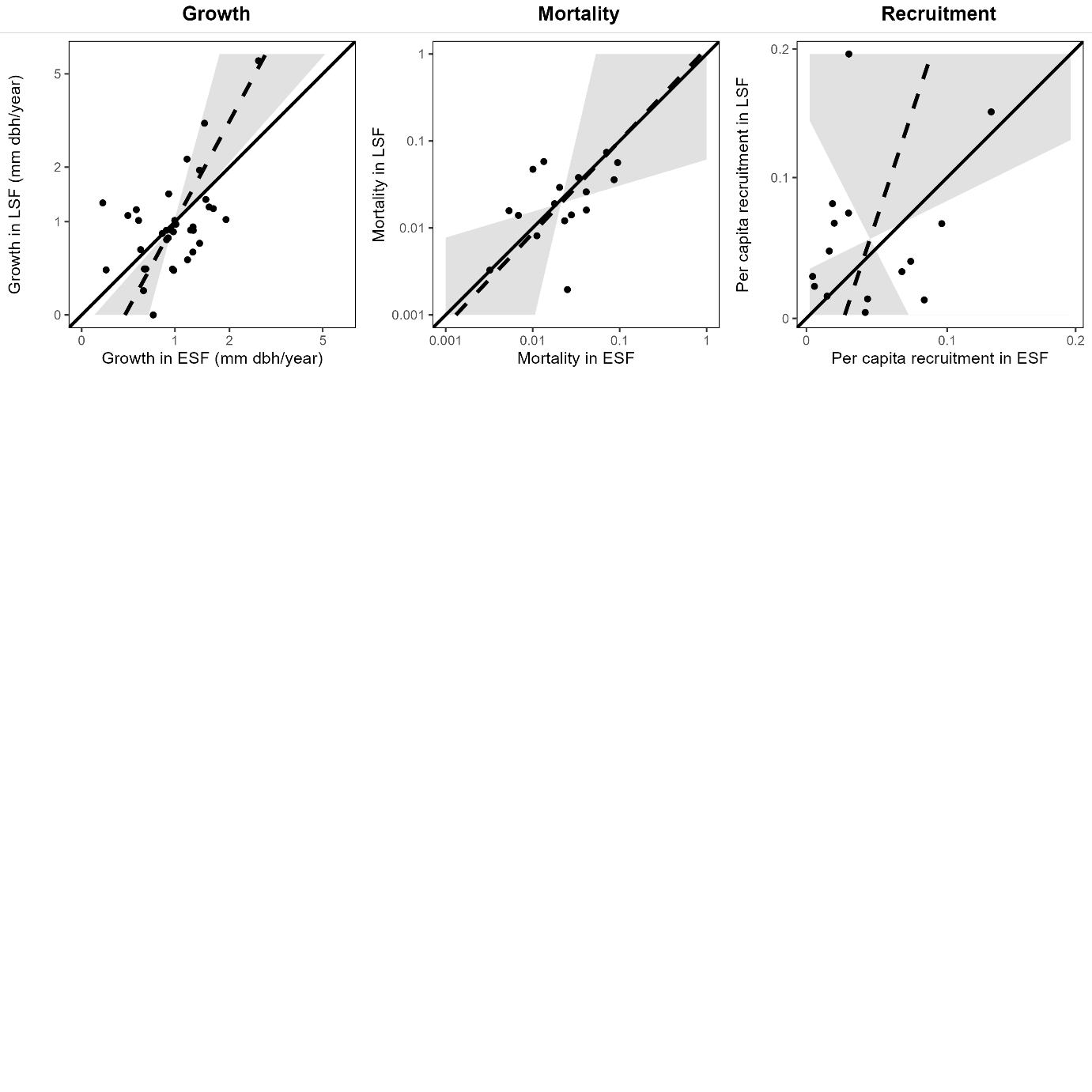

**Figure S13**: Major axis regressions for species’ demographic rates in early successional forests (ESF), late successional forests (LSF) and old-growth forests (OGF) in **Yucatán**. Each point represents a species and its demographic rate in the respective successional stage. Solid lines represent the 1:1-line, dashed lines represent the regression lines and areas highlighted in grey represent the confidence intervals. Growth and mortality rates are from canopy layer 2. Only species with at least 5 observations for growth and survival in both successional stages were included, respectively.

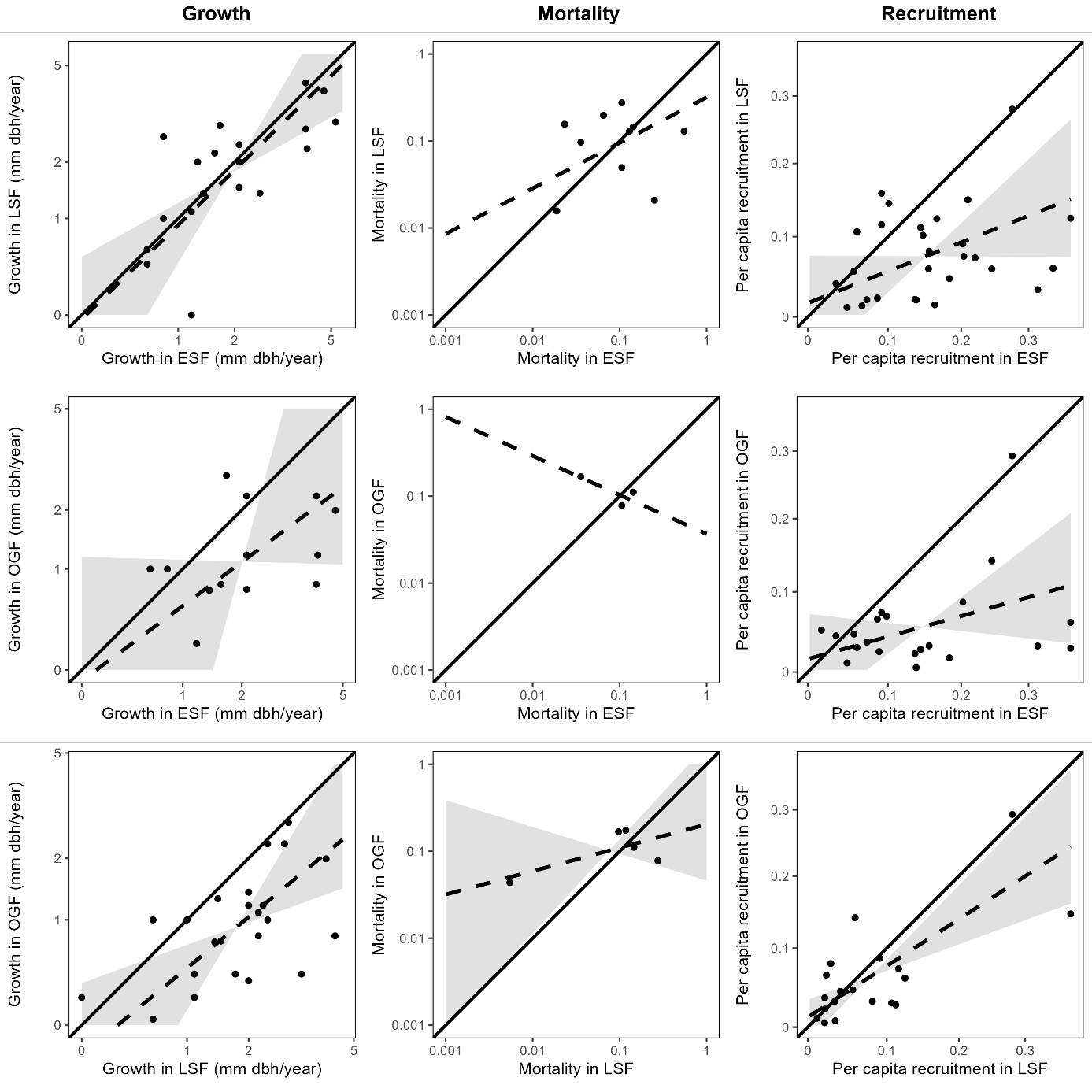

**Figure S14**: Major axis regressions for species’ demographic rates in early successional forests (ESF), late successional forests (LSF) and old-growth forests (OGF) in **Oaxaca**. Each point represents a species and its demographic rate in the respective successional stage. Solid lines represent the 1:1-line, dashed lines represent the regression lines and areas highlighted in grey represent the confidence intervals. Growth and mortality rates are from canopy layer 2. Only species with at least 5 observations for growth and survival in both successional stages were included, respectively. If no confidence intervals are given, the model was not statistically significant (i.e., the variables are unrelated).

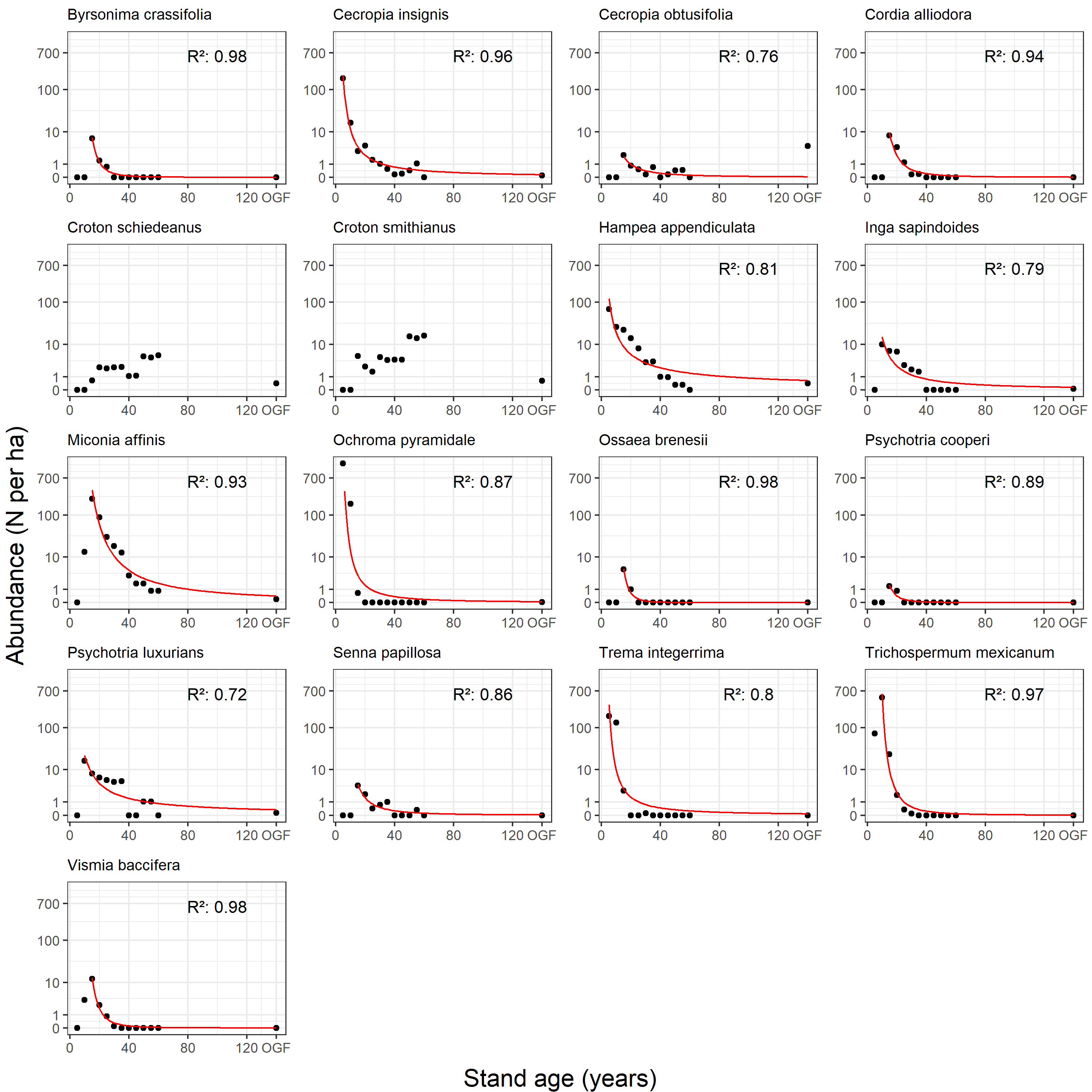

**Figure S15**: Observed (points) and modelled (red lines) abundances over time of individual species within the high mortality group exclusive to wet early successional forests in Costa Rica along with R² values for the models. Models are of the form Ln(abundance) = a*stand age^b. Parameters a and b were estimated using the *nls* R function. For model parameter estimation, we only used data points from the highest abundance onwards.

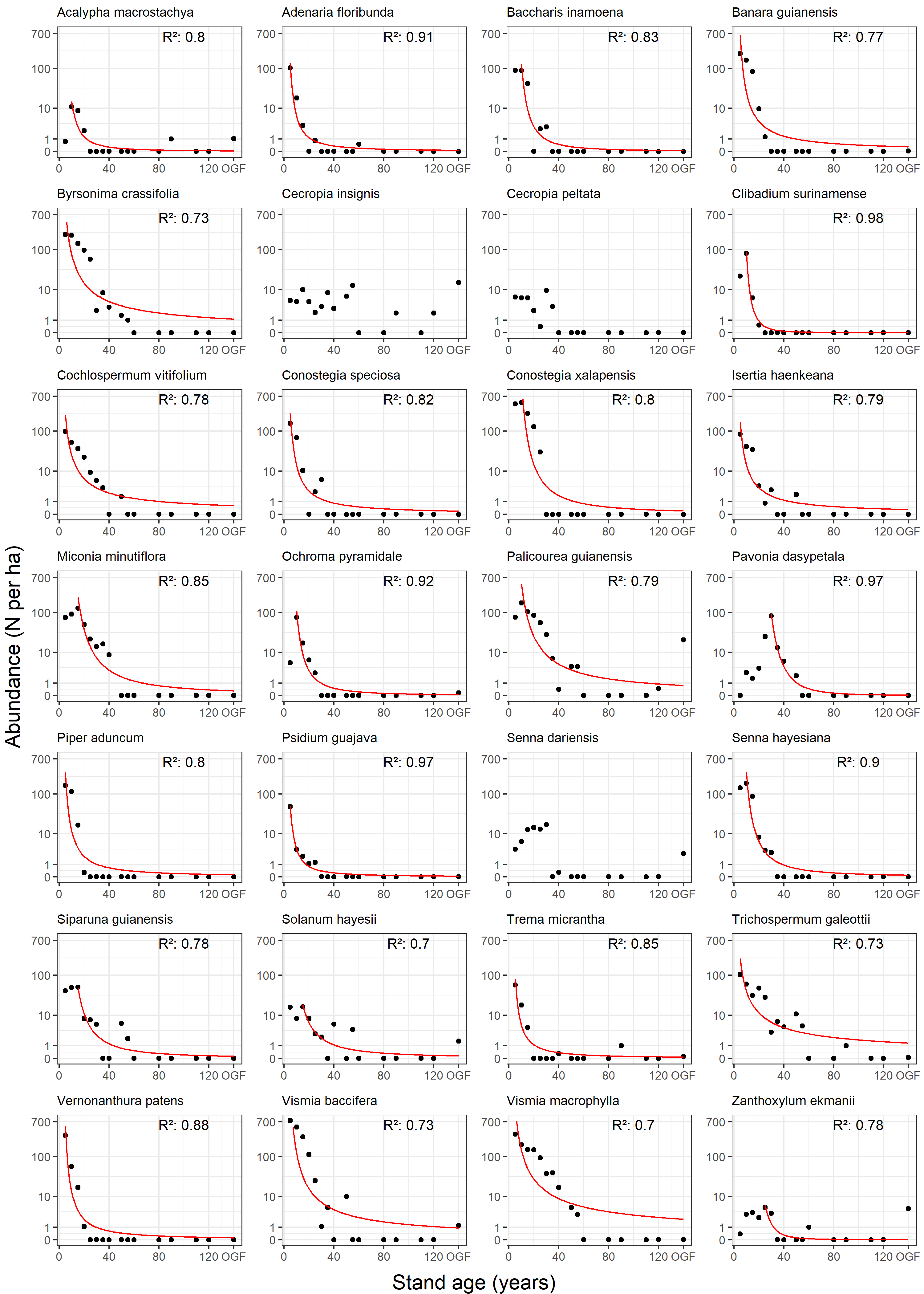

**Figure S16**: Observed (points) and modelled (red lines) abundances over time of individual species within the high mortality group exclusive to wet early successional forests in Panama along with R² values for the models. Models are of the form Ln(abundance) = a*stand age^b. Parameters a and b were estimated using the *nls* R function. For model parameter estimation, we only used data points from the highest abundance onwards.

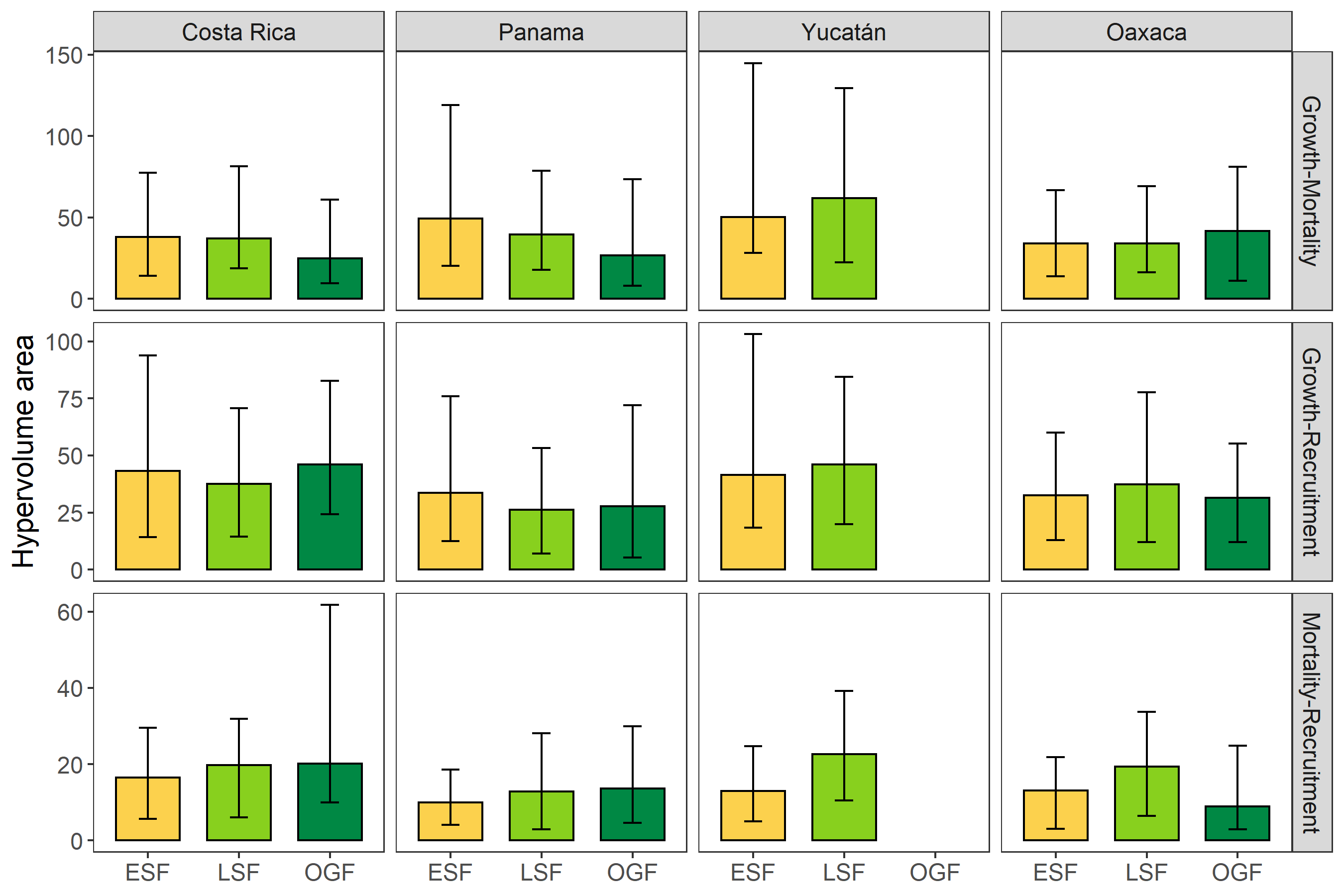

**Figure S17**: Areas of the two-dimensional hypervolumes representing the ranges of demographic strategies for different pairs of demographic rates and successional stages (ESF = early successional forest, LSF = late successional forest, OGF = old-growth forest). Colored bars represent the median rarefied and bootstrapped values, error bars represent 95% confidence intervals (r = 100 replicates, n = 10 species).

Poorter, L., Craven, D., Jakovac, C. C., van der Sande, M. T., Amissah, L., Bongers, F., Chazdon, R. L., Farrior, C. E., Kambach, S., Meave, J. A., Muñoz, R., Norden, N., Rüger, N., van Breugel, M., Almeyda Zambrano, A. M., Amani, B., Andrade, J. L., Brancalion, P. H. S., Broadbent, E. N., Foresta, H. de, Dent, D. H., Derroire, G., DeWalt, S. J., Dupuy, J. M., Durán, S. M., Fantini, A. C., Finegan, B., Hernández-Jaramillo, A., Hernández-Stefanoni, J. L., Hietz, P., Junqueira, A. B., N'dja, J. K., Letcher, S. G., Lohbeck, M., López-Camacho, R., Martínez-Ramos, M., Melo, F. P. L., Mora, F., Müller, S. C., N'Guessan, A. E., Oberleitner, F., Ortiz-Malavassi, E., Pérez-García, E. A., Pinho, B. X., Piotto, D., Powers, J. S., Rodríguez-Buriticá, S., Rozendaal, D. M. A., Ruíz, J., Tabarelli, M., Teixeira, H. M., Valadares de Sá Barretto Sampaio, E., van der Wal, H., Villa, P. M., Fernandes, G. W., Santos, B. A., Aguilar-Cano, J., Almeida-Cortez, J. S. de, Alvarez-Davila, E., Arreola-Villa, F., Balvanera, P., Becknell, J. M., Cabral, G. A. L., Castellanos-Castro, C., Jong, B. H. J. de, Nieto, J. E., Espírito-Santo, M. M., Fandino, M. C., García, H., García-Villalobos, D., Hall, J. S., Idárraga, A., Jiménez-Montoya, J., Kennard, D., Marín-Spiotta, E., Mesquita, R., Nunes, Y. R. F., Ochoa-Gaona, S., Peña-Claros, M., Pérez-Cárdenas, N., Rodríguez-Velázquez, J., Villanueva, L. S., Schwartz, N. B., Steininger, M. K., Veloso, M. D. M., Vester, H. F. M., Vieira, I. C. G., Williamson, G. B., Zanini, K., & Hérault, B. 2021. Multidimensional tropical forest recovery*.* Science 374: 1370–1376. https://doi.org/10.1126/science.abh3629.

Poorter, L., Kitajima, K., Mercado, P., Chubiña, J., Melgar, I., & Prins, H. H. T. 2010. Resprouting as a persistence strategy of tropical forest trees: relations with carbohydrate storage and shade tolerance*.* Ecology 91: 2613–2627. https://doi.org/10.1890/09-0862.1.

Poorter, L., Rozendaal, D. M. A., Bongers, F., Almeida-Cortez, J. S. de, Almeyda Zambrano, A. M., Álvarez, F. S., Andrade, J. L., Villa, L. F. A., Balvanera, P., Becknell, J. M., Bentos, T. V., Bhaskar, R., Boukili, V., Brancalion, P. H. S., Broadbent, E. N., César, R. G., Chave, J., Chazdon, R. L., Colletta, G. D., Craven, D., Jong, B. H. J. de, Denslow, J. S., Dent, D. H., DeWalt, S. J., García, E. D., Dupuy, J. M., Durán, S. M., Espírito Santo, M. M., Fandiño, M. C., Fernandes, G. W., Finegan, B., Moser, V. G., Hall, J. S., Hernández-Stefanoni, J. L., Jakovac, C. C., Junqueira, A. B., Kennard, D., Lebrija-Trejos, E., Letcher, S. G., Lohbeck, M., Lopez, O. R., Marín-Spiotta, E., Martínez-Ramos, M., Martins, S. V., Massoca, P. E. S., Meave, J. A., Mesquita, R., Mora, F., Souza Moreno, V. de, Müller, S. C., Muñoz, R., Muscarella, R., Oliveira Neto, S. N. de, Nunes, Y. R. F., Ochoa-Gaona, S., Paz, H., Peña-Claros, M., Piotto, D., Ruíz, J., Sanaphre-Villanueva, L., Sanchez-Azofeifa, A., Schwartz, N. B., Steininger, M. K., Thomas, W. W., Toledo, M., Uriarte, M., Utrera, L. P., van Breugel, M., van der Sande, M. T., van der Wal, H., Veloso, M. D. M., Vester, H. F. M., Vieira, I. C. G., Villa, P. M., Williamson, G. B., Wright, S. J., Zanini, K. J., Zimmerman, J. K., & Westoby, M. 2019. Wet and dry tropical forests show opposite successional pathways in wood density but converge over time*.* Nature Ecology & Evolution 3: 928–934. https://doi.org/10.1038/s41559-019-0882-6.

Poorter, L., van de Plassche, M., Willems, S., & Boot, R. G. A. 2004. Leaf traits and herbivory rates of tropical tree species differing in successional status*.* Plant Biology 6: 746–754. https://doi.org/10.1055/s-2004-821269.

Purves, D. W., Lichstein, J. W., Strigul, N., & Pacala, S. W. 2008. Predicting and understanding forest dynamics using a simple tractable model*.* Proceedings of the National Academy of Sciences of the United States of America 105: 17018–17022. https://doi.org/10.1073/pnas.0807754105.

R Core Team. 2022. R: A language and environment for statistical computing. R Foundation for Statistical Computing, Vienna, Austria.

Rico-Gray, V., & García-Franco, J. G. 1991. The maya and the vegetation of the Yucatán peninsula*.* Journal of Ethnobiology 11: 135–142.

Rozendaal, D. M. A., Bongers, F., Aide, T. M., Alvarez-Dávila, E., Ascarrunz, N., Balvanera, P., Becknell, J. M., Bentos, T. V., Brancalion, P. H. S., Cabral, G. A. L., Calvo-Rodriguez, S., Chave, J., César, R. G., Chazdon, R. L., Condit, R., Dallinga, J. S., Almeida-Cortez, J. S. de, Jong, B. de, Oliveira, A. de, Denslow, J. S., Dent, D. H., DeWalt, S. J., Dupuy, J. M., Durán, S. M., Dutrieux, L. P., Espírito-Santo, M. M., Fandino, M. C., Fernandes, G. W., Finegan, B., García, H., Gonzalez, N., Moser, V. G., Hall, J. S., Hernández-Stefanoni, J. L., Hubbell, S., Jakovac, C. C., Hernández, A. J., Junqueira, A. B., Kennard, D., Larpin, D., Letcher, S. G., Licona, J.-C., Lebrija-Trejos, E., Marín-Spiotta, E., Martínez-Ramos, M., Massoca, P. E. S., Meave, J. A., Mesquita, R. C. G., Mora, F., Müller, S. C., Muñoz, R., Oliveira Neto, S. N. de, Norden, N., Nunes, Y. R. F., Ochoa-Gaona, S., Ortiz-Malavassi, E., Ostertag, R., Peña-Claros, M., Pérez-García, E. A., Piotto, D., Powers, J. S., Aguilar-Cano, J., Rodriguez-Buritica, S., Rodríguez-Velázquez, J., Romero-Romero, M. A., Ruíz, J., Sanchez-Azofeifa, A., Almeida, A. S. de, Silver, W. L., Schwartz, N. B., Thomas, W. W., Toledo, M., Uriarte, M., Sá Sampaio, E. V. de, van Breugel, M., van der Wal, H., Martins, S. V., Veloso, M. D. M., Vester, H. F. M., Vicentini, A., Vieira, I. C. G., Villa, P., Williamson, G. B., Zanini, K. J., Zimmerman, J., & Poorter, L. 2019. Biodiversity recovery of Neotropical secondary forests*.* Science Advances 5: eaau3114. https://doi.org/10.1126/sciadv.aau3114.

Rüger, N., Comita, L. S., Condit, R., Purves, D., Rosenbaum, B., Visser, M. D., Wright, S. J., & Wirth, C. 2018. Beyond the fast-slow continuum: demographic dimensions structuring a tropical tree community*.* Ecology Letters 21: 1075–1084. https://doi.org/10.1111/ele.12974.

Rüger, N., Condit, R., Dent, D. H., DeWalt, S. J., Hubbell, S. P., Lichstein, J. W., Lopez, O. R., Wirth, C., & Farrior, C. E. 2020. Demographic trade-offs predict tropical forest dynamics*.* Science 368: 165–168. https://doi.org/10.1126/science.aaz4797.

Rüger, N., Schorn, M. E., Kambach, S., Chazdon, R. L., Farrior, C. E., Meave, J. A., Muñoz, R., van Breugel, M., Amissah, L., Bongers, F., Craven, D., Hérault, B., Jakovac, C. C., Norden, N., Poorter, L., van der Sande, M. T., Wirth, C., Delgado, D., Dent, D. H., DeWalt, S. J., Dupuy, J. M., Finegan, B., Hall, J. S., Hernandez-Stefanoni, J. L., & Lopez, O. R. submitted. Successional shifts in tree demographic strategies in Neotropical wet and dry forests.

Rüger, N., Wirth, C., Wright, S. J., & Condit, R. 2012. Functional traits explain light and size response of growth rates in tropical tree species*.* Ecology 93: 2626–2636. https://doi.org/10.1890/12-0622.1.

Russo, S. E., McMahon, S. M., Detto, M., Ledder, G., Wright, S. J., Condit, R. S., Davies, S. J., Ashton, P. S., Bunyavejchewin, S., Chang-Yang, C.-H., Ediriweera, S., Ewango, C. E. N., Fletcher, C., Foster, R. B., Gunatilleke, C. V. S., Gunatilleke, I. A. U. N., Hart, T., Hsieh, C.-F., Hubbell, S. P., Itoh, A., Kassim, A. R., Leong, Y. T., Lin, Y. C., Makana, J.-R., Mohamad, M. B., Ong, P., Sugiyama, A., Sun, I.-F., Tan, S., Thompson, J., Yamakura, T., Yap, S. L., & Zimmerman, J. K. 2021. The interspecific growth-mortality trade-off is not a general framework for tropical forest community structure*.* Nature Ecology & Evolution 5: 174–183. https://doi.org/10.1038/s41559-020-01340-9.

Saenz-Pedroza, I., Feldman, R., Reyes-García, C., Meave, J. A., Calvo-Irabien, L. M., May-Pat, F., & Dupuy, J. M. 2020. Seasonal and successional dynamics of size-dependent plant demographic rates in a tropical dry forest*.* PeerJ 8: e9636. https://doi.org/10.7717/peerj.9636.

Salguero-Gómez, R., Jones, O. R., Jongejans, E., Blomberg, S. P., Hodgson, D. J., Mbeau-Ache, C., Zuidema, P. A., Kroon, H. de, & Buckley, Y. M. 2016. Fast-slow continuum and reproductive strategies structure plant life-history variation worldwide*.* Proceedings of the National Academy of Sciences of the United States of America 113: 230–235. https://doi.org/10.1073/pnas.1506215112.

Sanaphre-Villanueva, L., Dupuy, J. M., Andrade, J. L., Reyes-García, C., Jackson, P. C., & Paz, H. 2017. Patterns of plant functional variation and specialization along secondary succession and topography in a tropical dry forest*.* Environmental Research Letters 12: 55004. https://doi.org/10.1088/1748-9326/aa6baa.

Schnitzer, S. A., & Carson, W. P. 2001. Treefall gaps and the maintenance of species diversity in a tropical forest*.* Ecology 82: 913–919. https://doi.org/10.1890/0012-9658(2001)082[0913:TGATMO]2.0.CO;2.

Slik, J. W. F., Arroyo-Rodríguez, V., Aiba, S.-I., Alvarez-Loayza, P., Alves, L. F., Ashton, P., Balvanera, P., Bastian, M. L., Bellingham, P. J., van den Berg, E., Bernacci, L., Da Conceição Bispo, P., Blanc, L., Böhning-Gaese, K., Boeckx, P., Bongers, F., Boyle, B., Bradford, M., Brearley, F. Q., Breuer-Ndoundou Hockemba, M., Bunyavejchewin, S., Calderado Leal Matos, D., Castillo-Santiago, M., Catharino, E. L. M., Chai, S.-L., Chen, Y., Colwell, R. K., Chazdon, R. L., Robin, C. L., Clark, C., Clark, D. B., Clark, D. A., Culmsee, H., Damas, K., Dattaraja, H. S., Dauby, G., Davidar, P., DeWalt, S. J., Doucet, J.-L., Duque, A., Durigan, G., Eichhorn, K. A. O., Eisenlohr, P. V., Eler, E., Ewango, C., Farwig, N., Feeley, K. J., Ferreira, L., Field, R., Oliveira Filho, A. T. de, Fletcher, C., Forshed, O., Franco, G., Fredriksson, G., Gillespie, T., Gillet, J.-F., Amarnath, G., Griffith, D. M., Grogan, J., Gunatilleke, N., Harris, D., Harrison, R., Hector, A., Homeier, J., Imai, N., Itoh, A., Jansen, P. A., Joly, C. A., Jong, B. H. J. de, Kartawinata, K., Kearsley, E., Kelly, D. L., Kenfack, D., Kessler, M., Kitayama, K., Kooyman, R., Larney, E., Laumonier, Y., Laurance, S., Laurance, W. F., Lawes, M. J., Amaral, I. L. d., Letcher, S. G., Lindsell, J., Lu, X., Mansor, A., Marjokorpi, A., Martin, E. H., Meilby, H., Melo, F. P. L., Metcalfe, D. J., Medjibe, V. P., Metzger, J. P., Millet, J., Mohandass, D., Montero, J. C., Morisson Valeriano, M. de, Mugerwa, B., Nagamasu, H., Nilus, R., Ochoa-Gaona, S., Onrizal, Page, N., Parolin, P., Parren, M., Parthasarathy, N., Paudel, E., Permana, A., Piedade, M. T. F., Pitman, N. C. A., Poorter, L., Poulsen, A. D., Poulsen, J., Powers, J., Prasad, R. C., Puyravaud, J.-P., Razafimahaimodison, J.-C., Reitsma, J., Dos Santos, J. R., Roberto Spironello, W., Romero-Saltos, H., Rovero, F., Rozak, A. H., Ruokolainen, K., Rutishauser, E., Saiter, F., Saner, P., Santos, B. A., Santos, F., Sarker, S. K., Satdichanh, M., Schmitt, C. B., Schöngart, J., Schulze, M., Suganuma, M. S., Sheil, D., Da Silva Pinheiro, E., Sist, P., Stevart, T., Sukumar, R., Sun, I.-F., Sunderland, T., Sunderand, T., Suresh, H. S., Suzuki, E., Tabarelli, M., Tang, J., Targhetta, N., Theilade, I., Thomas, D. W., Tchouto, P., Hurtado, J., Valencia, R., van Valkenburg, J. L. C. H., van Do, T., Vasquez, R., Verbeeck, H., Adekunle, V., Vieira, S. A., Webb, C. O., Whitfeld, T., Wich, S. A., Williams, J., Wittmann, F., Wöll, H., Yang, X., Adou Yao, C. Y., Yap, S. L., Yoneda, T., Zahawi, R. A., Zakaria, R., Zang, R., Assis, R. L. de, Garcia Luize, B., & Venticinque, E. M. 2015. An estimate of the number of tropical tree species*.* Proceedings of the National Academy of Sciences of the United States of America 112: 7472–7477. https://doi.org/10.1073/pnas.1423147112.

Stearns, S. C. 1992. The evolution of life histories. Oxford University Press, Oxford.

Tilman, D. 1988. Plant strategies and the dynamics and structure of plant communities. Princeton Univ. Press, Princeton, NJ.

van Breugel, M., Hall, J. S., Craven, D., Bailon, M., Hernandez, A., Abbene, M., & van Breugel, P. 2013. Succession of ephemeral secondary forests and their limited role for the conservation of floristic diversity in a human-modified tropical landscape*.* PloS one 8: e82433. https://doi.org/10.1371/journal.pone.0082433.

Vieira, D. L. M., & Scariot, A. 2006. Principles of natural regeneration of tropical dry forests for restoration*.* Restoration Ecology 14: 11–20. https://doi.org/10.1111/j.1526-100X.2006.00100.x.

Walker, L. R., Wardle, D. A., Bardgett, R. D., & Clarkson, B. D. 2010. The use of chronosequences in studies of ecological succession and soil development*.* Journal of Ecology 98: 725–736. https://doi.org/10.1111/j.1365-2745.2010.01664.x.

Wright, S. J., Kitajima, K., Kraft, N. J. B., Reich, P. B., Wright, I. J., Bunker, D. E., Condit, R., Dalling, J. W., Davies, S. J., Díaz, S., Engelbrecht, B. M. J., Harms, K. E., Hubbell, S. P., Marks, C. O., Ruiz-Jaen, M. C., Salvador, C. M., & Zanne, A. E. 2010. Functional traits and the growth-mortality trade-off in tropical trees*.* Ecology 91: 3664–3674. https://doi.org/10.1890/09-2335.1.
